# High Cysteinyl Leukotriene Receptor 1 Expression Correlates with Poor Survival of Uveal Melanoma Patients and Cognate Antagonist Drugs Modulate the Growth, Cancer Secretome, and Metabolism of Uveal Melanoma Cells

**DOI:** 10.1101/2020.08.23.261545

**Authors:** Kayleigh Slater, Aisling B. Heeran, Rebeca Sanz-Pamplona, Helen Kalirai, Arman Rahman, Mays Helmi, Sandra Garcia-Mulero, Fiona O’Connell, Rosa Bosch, Anna Portela, Alberto Villanueva, Josep M. Piulats, William M. Gallagher, Lasse D. Jensen, Sarah E. Coupland, Jacintha O’Sullivan, Breandán N. Kennedy

## Abstract

Uveal melanoma (UM) is a rare, but often lethal, form of ocular cancer arising from melanocytes within the uveal tract. UM has a high propensity to spread hematogenously to the liver, with up to 50% of patients developing liver metastases. Unfortunately, once liver metastasis occurs, patient prognosis is extremely poor with as few as 8% of patients surviving beyond two years. There are no *standard-of-care* therapies available for the treatment of metastatic uveal melanoma, hence it is a clinical area of urgent unmet need. Here, the clinical relevance and therapeutic potential of cysteinyl leukotriene receptors (CysLT_1_ and CysLT_2_) in UM was evaluated. High expression of *CYSLTR1* or *CYSLTR2* transcripts is significantly associated with poor disease-free survival and poor overall survival in UM patients. Digital pathology analysis identified high expression of CysLT_1_ in primary UM is associated with reduced disease-specific survival (p = 0.012) and overall survival (p = 0.011). High CysLT_1_ expression shows a statistically significant (p = 0.041) correlation with ciliary body involvement, a poor prognostic indicator in UM. Small molecule drugs targeting CysLT_1_ were vastly superior at exerting anti-cancer phenotypes in UM cell lines and zebrafish xenografts than drugs targeting CysLT_2_. Quininib, a selective CysLT_1_ antagonist, significantly inhibits survival (p < 0.0001), long-term proliferation (p < 0.0001), and oxidative phosphorylation (p < 0.001), but not glycolysis, in primary and metastatic UM cell lines. Quininib exerts opposing effects on the secretion of inflammatory markers in primary versus metastatic UM cell lines. Quininib significantly downregulated IL-2 and IL-6 in Mel285 cells (p < 0.05), but significantly upregulated IL-10, IL-1β, IL-2 (p < 0.0001), IL-13, IL-8 (p < 0.001), IL-12p70 and IL-6 (p < 0.05) in OMM2.5 cells. Finally, quininib significantly inhibits tumour growth in orthotopic zebrafish xenograft models of UM. These preclinical data suggest that antagonism of CysLT_1_, but not CysLT_2_, may be of therapeutic interest in the treatment of UM.

## 1. Introduction

Uveal melanoma (UM) is a rare, intraocular cancer that metastasises predominantly to the liver in approximately 50% of patients. The primary ocular tumour is usually successfully treated with surgery or radiotherapy [1,2]. However, these treatments have limited success in halting metastatic spread of the disease. Once the cancer has disseminated to the liver, there are limited options available to patients. Overall survival for patients with metastatic UM ranges from 4 to 18 months [3-6]. UM develops in one of the most capillary-rich tissues of the body and is spread solely through the blood stream, suggesting that angiogenesis and vascular invasion play important roles in UM progression.

In comparison to other solid tumours and skin melanomas, UM has a low mutational burden [7,8] making the disease less sensitive to checkpoint inhibitors. The lack of identified mutations in UM narrows the scope for targeted therapies, with no successful targeted therapies available to date [9]. Activating mutations in *GNAQ* or *GNA11* are found in >80% of all UMs [10], with mutations in *CYSLTR2* or *PLCB4* likely to account for an additional 8-10% of activating UM mutations [11]. These mutations are mutually exclusive and operate in the same pathway [12], highlighting the importance of CysLT_2_/G_αq/11_/PLCB4 signalling in UM oncogenesis. In contrast to cutaneous melanoma [13], targeted therapies for UM, including those targeting the CysLT_2_/G_αq/11_/PLCB4 downstream pathways, such as MEK and AKT, failed in early clinical studies [14,15].

Synthesised through the 5-lipoxygenase (5-LO) pathway, the cysteinyl leukotrienes (CysLTs), LTC_4_, LTD_4_ and LTE_4_, are lipid-signalling molecules that mediate acute and chronic inflammation [16]. The CysLTs exert their biological effects via binding to the G protein-coupled receptors (GPCRs), CysLT_1_ and CysLT_2_. LTD_4_ binds to CysLT_1_ with high affinity [17], while both LTD_4_ and LTC_4_ bind to CysLT_2_ with equal affinity [18]. Although activation of both receptors stimulates similar downstream signalling events (calcium flux and accumulation of inositol phosphate) [17,18], the receptors are not functionally redundant [19]. Each receptor has a distinct pattern of cellular and tissue expression [17,18], which in combination with their differing sensitivities to endogenous leukotriene ligands, suggests each receptor has an individual role in physiology and pathology [20]. Cross-regulation occurs between the receptors: CysLT_2_ controls the membrane expression of CysLT_1_ and negatively regulates signalling through CysLT_1_ [19].

CysLTs are well known for their role in inflammation, particularly in asthma and allergic rhinitis. Recently, however, a role for CysLTs in cancer has emerged [9,21], with a particular focus on their role in vascular permeability and angiogenesis [22]. In a retrospective analysis, CysLT_1_ antagonists, montelukast and zafirlukast, display a dose-dependent chemopreventative effect against 14 different cancers [23]. Furthermore, overexpression of CysLT_1_ is a feature of colorectal cancer, prostate cancer, renal cell carcinoma, urothelial transitional cell carcinoma and testicular cancer [24-27]. Interestingly, colorectal and breast cancer patients with high expression of CysLT_1_ have a poor prognosis and reduced survival, respectively [28,29]. In contrast, a recurrent, hotspot mutation in *CYSLTR2* is a driver oncogene in a small subset of UM [12]. This mutation encodes a p.Leu129Gln substitution, which leads to constitutive activation of endogenous G_αq_ signalling and promotes tumorigenesis *in vivo* [12]. The same Leu129Gln hotspot mutation in *CYSLTR2* has also been identified in blue nevi [30], and in leptomeningeal melanocytic tumours [31] confirming the oncogenic properties of constitutively active signalling through this receptor.

Quininib, a CysLT_1_ antagonist, and its analogue 1,4-dihydroxy quininib, are anti-inflammatory and anti-angiogenic drugs [32,33] with anti-cancer activity in human *ex vivo* colorectal cancer patient tumour explants and colorectal cancer xenograft models [34,35]. Angiogenesis plays an important role in the development and progression of UM, however the efficacy of anti-angiogenic therapies remains inadequate [36]. In addition, highly vascularised UM are more aggressive and convey a worse prognosis [37,38]. This, coupled with the finding that *CYSLTR2* acts as an oncogene in a subset of UM, led us to hypothesise that CysLT receptors represent a therapeutic target for UM. To our knowledge, this is the first study to examine the clinical significance of CysLT receptor gene and protein expression in UM patients. Critically, we find that high expression of *CYSLTR1* and *CYSLTR2* transcripts correlate with reduced disease-free survival and reduced overall survival in UM patients. We show that both CysLT receptors are expressed in primary UM and that high expression of CysLT_1_ is linked to ciliary body involvement, a poor prognostic indicator in UM. Additionally, high expression of CysLT_1_ in primary UM is significantly associated with reduced melanoma-specific survival and overall survival. CysLT_1_ antagonists alter survival, long-term proliferation, metabolism, and the secretion of proinflammatory and proangiogenic factors in *in vitro* UM cell line models, and can promote anti-cancer activity in *in vivo* zebrafish xenograft models of UM. This study reinforces the importance of the *CysLT*_*1/2*_ pathway in UM and suggests that inhibition of CysLT_1_ represents an alternative therapeutic target for UM.

## 2. Results

### 2.1. Analysis of cysteinyl leukotriene receptor expression in primary uveal melanoma patient samples

The association between the expression of CysLT receptors in primary UM samples and the associated clinical features has not been previously reported. We examined whether CysLT receptors were expressed in UM patient samples and whether their expression correlated with clinical outcomes.

The Cancer Genome Atlas (TCGA) contains gene expression data on 80 primary UM samples. Both CysLT receptor genes were expressed in the UM samples confirming their potential disease relevance. Cox survival analysis revealed expression of CYSLTR1 and CYSLTR2 genes are significantly associated with disease free survival (p = 0.014 and p = 0.003, respectively) (Figure 1A and D) and overall survival (p = 0.004 and p = 0.0008, respectively)(Figure 1B and E) in UM patients. When stratified based on high versus low expression of the receptors, all Kaplan-Meier curves showed significant results. This suggests that UM patients with high CYSLTR1 or CYSLTR2 transcript expression have a worse prognosis than those with low expression of the receptors. TCGA-UM samples were then divided into high and low expression of CYSLTR1 and CYSLTR2 using the third quartile as cut-off and interrogated for their association with pathways of interest. Samples expressing higher CYSLTR1 showed significant overexpression of Inflammatory Response, IFN-γ, TNF-α, Angiogenesis and GPCR signalling (Figure 1C). Interestingly, the same pathways plus glycolysis were overexpressed in samples with high expression of CYSLTR2 (Figure 1F).

**Figure 1.**
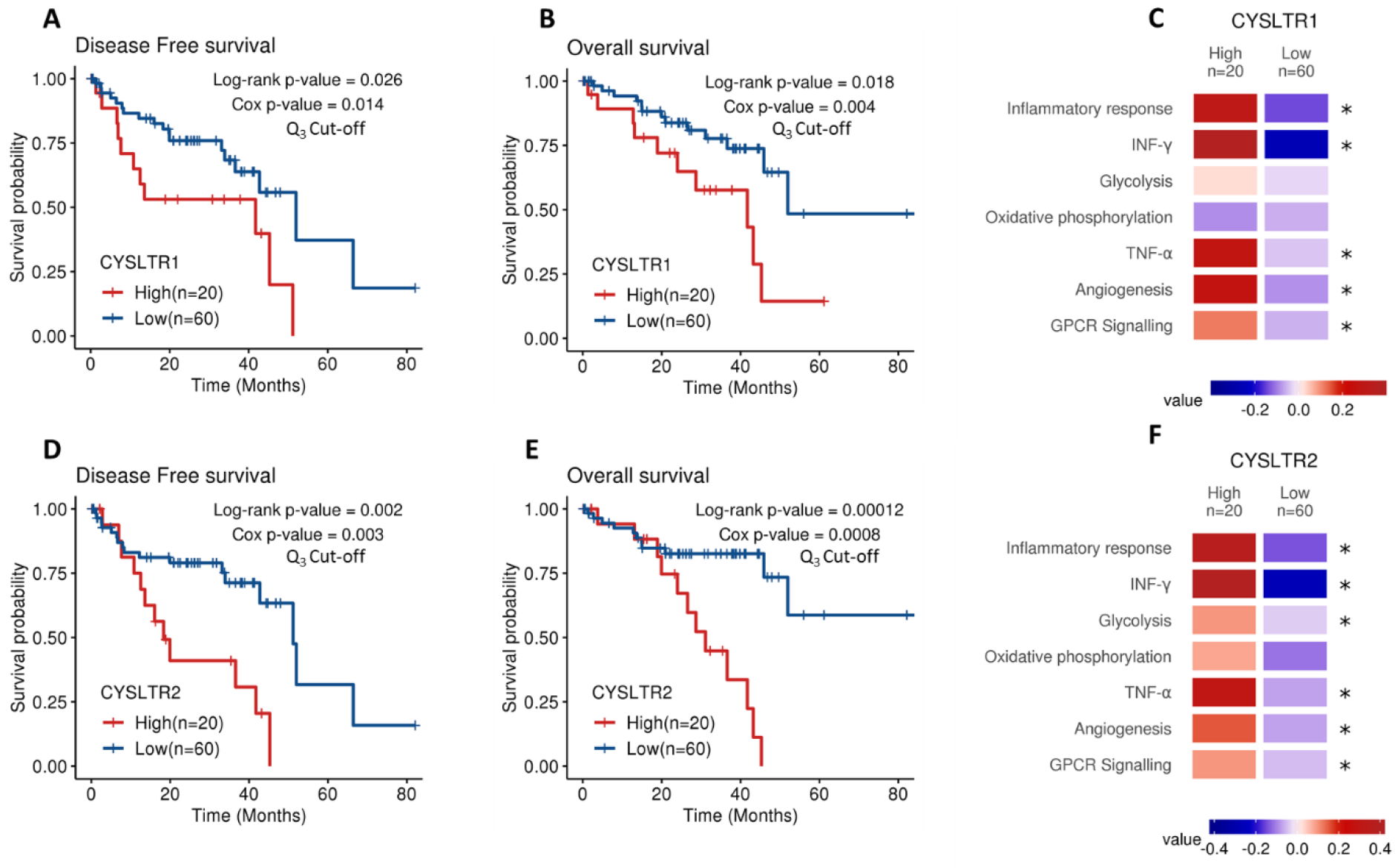
Analysis of *CYSLTR1/CYSLTR2* expression and UM patient survival from The Cancer Genome Atlas (TCGA).

Following analysis of gene expression profiles for both receptors, we examined the protein expression and localisation in an independent cohort of patients. The clinical relevance of CysLT receptors in UM was evaluated by analysing the expression of CysLT_1_ and CysLT_2_ proteins in a (TMA) generated from primary UM of 52 consented patients treated at the Liverpool Ocular Oncology Centre.

Of these 52 patients, 26 were males and 26 females, with a median age of 62 years at primary management (range, 39 −89). Survival data were not available for one patient, which reduced the numbers available for survival analysis to 51 patients. At the time of study end (29^th^ May 2019), 18/51 UM patients were alive (35.29%), 25/51 had died from metastatic disease (49.02%) and 8/25 died from other causes (15.69%). The median survival time was 8 years (range, 0.4 – 19 years). The median largest ultrasound diameter was 17.35 mm (range, 10.6 – 23.6 mm) with a median ultrasound tumour height of 8 mm (range, 5 – 18 mm)(Supplementary Figure 1C). Epithelioid cells were present in 30/52 (57.69%) of UM cases; 15/52 (28.85%) tumours involved the ciliary body and 3/52 (5.78%) had extraocular extension (Supplementary Figure 1D).

To conduct manual analysis for both CysLT_1_ and CysLT_2_, scores were assigned based on staining intensity (Figure 2A & D) and the percentage of tumour cells stained combined. Using the median as a cut off, a score of 0-7 was designated as low CysLT_1/2_ expression, while a score of 8-12 was designated as high CysLT_1/2_ expression. In Kaplan-Meier survival curves generated from median scoring of all primary UM cases, immunohistochemical levels of CysLT_1_ or CysLT_2_ did not demonstrate a significant association with patient survival from metastatic disease (CysLT_1_ p = 0.122, CysLT_2_ p = 0.341) (Figure 2B & E, respectively). However, high CysLT_1_ expression showed a robust trend towards reduced patient survival (Figure 2B). In agreement with the TCGA data, manual analysis of the UM TMA at the median cut-off revealed a statistically significant relationship between high CysLT_1_ expression and overall survival in the UM patients (p = 0.034) (Figure 2C). High CysLT_1_ expression also had a statistically significant relationship with ciliary body involvement in the UM patient cohort (p = 0.041) (Supplementary Figure 1D). Ciliary body involvement is a feature of the disease associated with metastatic risk and poor patient prognosis [39,40]. High expression of CysLT_2_ was not significantly associated with overall survival (p = 0.697) (Figure 2F). There was no statistically significant relationship between high or low CysLT_1_ expression and other clinical features of disease (extraocular extension, monosomy 3, periodic acid-Schiff positive loops, or epithelioid cell morphology) examined (Supplementary Figure 1D). Manual analysis did not identify a statistically significant relationship between high or low CysLT_2_ expression and clinical features of the disease examined (Supplementary Figure 1D).

**Figure 2.**
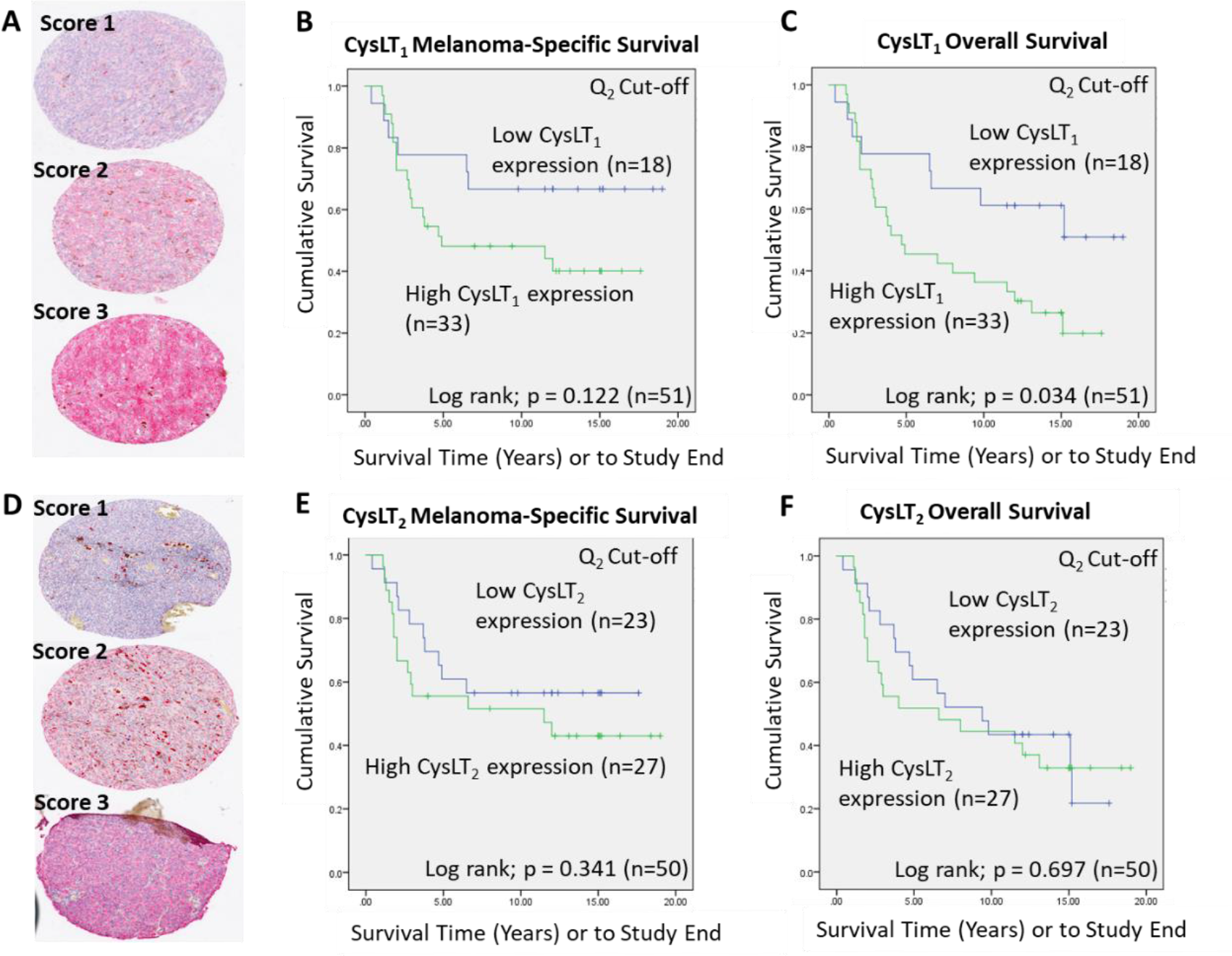
Examination of the prognostic value of CysLT_1_ and CysLT_2_ protein expression in primary UM samples by manual pathology.

A wider scoring range achieved by digital pathology analysis of this TMA strengthened the relationship between high CysLT_1_ expression and patient survival. With a 3^rd^ quartile segregation, high expression of CysLT_1_ is significantly associated with reduced melanoma-specific survival (p = 0.0012) and reduced overall survival (p = 0.0011) in this primary UM cohort (Figure 3B & C, respectively). In agreement with manual analysis, there was no significant relationship between CysLT_2_ expression and patient outcomes (Figure 3D & E).

**Figure 3.**
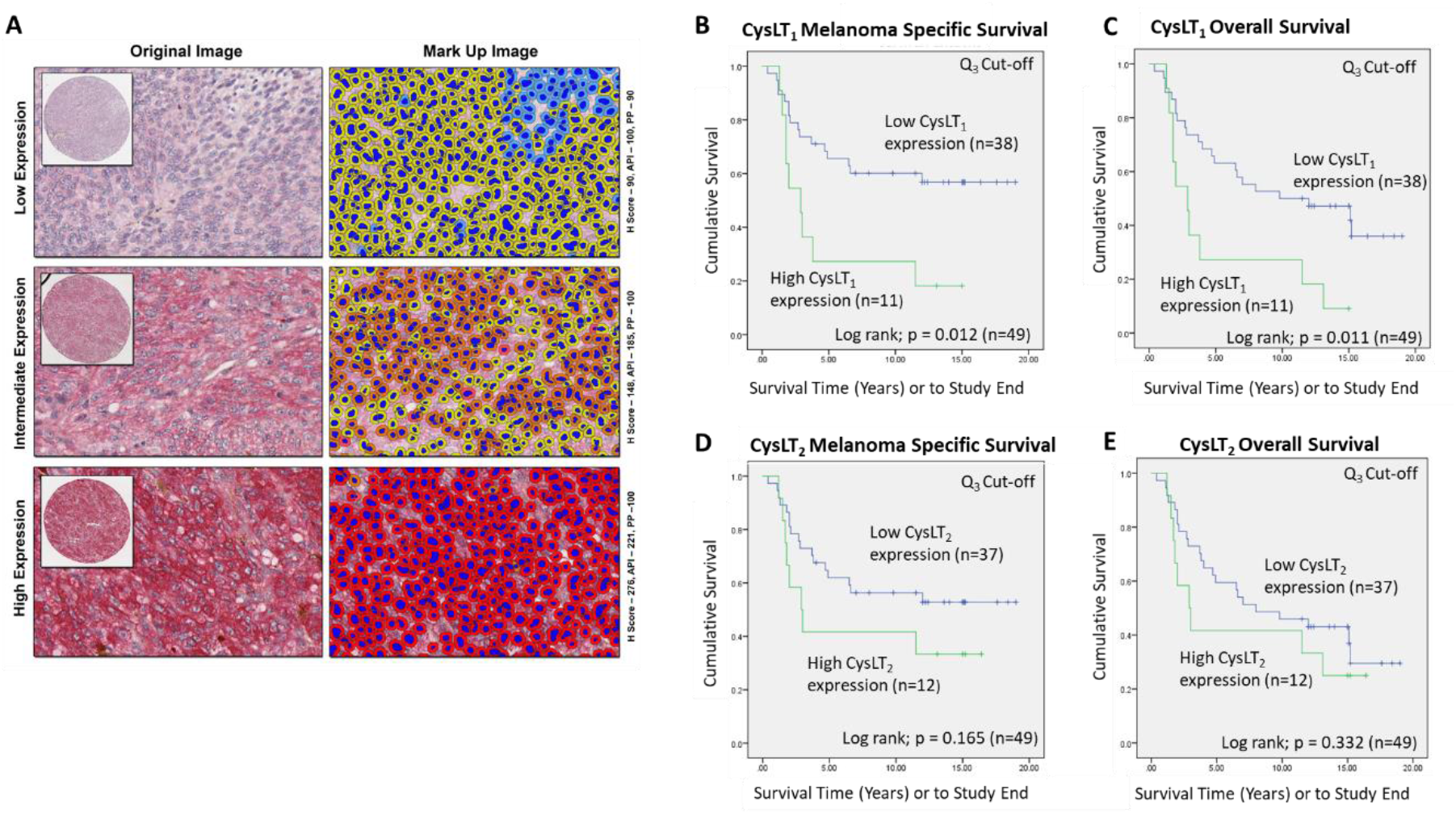
Examination of the prognostic value of CysLT_1_ and CysLT_2_ protein expression in primary UM patient samples by digital pathology analysis.

Gene expression data from TCGA suggests that high expression of CysLT_2_ is significantly linked to overall survival in UM patients. This was not supported by the data generated at the protein level using the UM patient TMA. Data from both TCGA and the UM patient TMA suggest a link between high CysLT_1_ expression and patient survival. A significant relationship between high CysLT_1_ expression and melanoma specific survival, as well as overall survival, was confirmed by our data. Interestingly, high expression of CysLT_1_ was significantly associated with ciliary body involvement, further suggesting a potential link between receptor expression and patient prognosis.

Kaplan-Meier survival curves demonstrate a statistically significant relationship between high (red) *CYSLTR1* expression and disease-free survival (A) (n =80; Log-rank; 0.026) or overall survival (B) (n =80; Log-rank; 0.018) in UM patients. Low *CYSLTR1* expression is shown in blue. Similarly, Kaplan-Meier survival curves demonstrate a statistically significant relationship between high (red) *CYSLTR2* expression and disease-free survival (D) (n =80; Log-rank; 0.002), or overall survival (E) (n =80; Log-rank; 0.00012) in UM patients. Low *CYSLTR2* expression is shown in blue. The third quartile was used as the cut-off point for high versus low expression for all Kaplan-Meier survival curves. Both Log-rank p-values (categorical variable) and Cox p-values (continuous variable) were calculated and are displayed.

Functional enrichment analysis: Samples were scored using gene expression profiles and categorized into high and low *CYSLTR1*(C) and *CYSLTR2* (F) expression using the third quartile as cut-off. Samples with high expression of *CYSLTR1* showed significant overexpression of an Inflammatory Response, INF-γ, TNF-α, Angiogenesis and GPCR signalling (C) (p < 0.05). Samples with high expression of *CYSLTR2* showed significant overexpression of Inflammatory Response, IFN-γ, Glycolysis, TNF-α, Angiogenesis and GPCR signalling (F) (p < 0.05).

(A) Representative cores from the UM patient tissue microarray (TMA) designated with a score of 1, 2 or 3 for CysLT_1_ staining intensity. (B) Kaplan-Meier survival curve stratified based by high (green) or low (blue) CysLT_1_ expression and death by metastatic melanoma (n=51; Log Rank; p =0.122). (C) High expression of CysLT_1_ (green) is significantly associated with reduced overall survival in primary UM patients (n =51; Log Rank; p= 0.034) (D) Representative cores from the TMA designated with a score of 1, 2 or 3 for CysLT_2_ staining intensity. (E). Kaplan-Meier survival curve stratified based by high (green) or low (blue) CysLT_2_ expression and death by metastatic melanoma (n=50; Log Rank; p =0.341). (F) Kaplan-Meier survival curve stratified based by high (green) or low (blue) CysLT_2_ expression and death by any cause (n=50; Log Rank; p =0.697). The median was used as the cut-off point for high versus low expression for all Kaplan-Meier survival curves. Number of events indicates the number of deaths due to metastatic melanoma (B,E). Number of events indicates the number of deaths due to any cause (C,F).

(A) Representative cores from the UM TMA designated with a score of Low, Intermediate or High expression for CysLT_1_ staining following digital analysis. (B) High expression of CysLT_1_ (green) is associated with reduced survival from metastatic melanoma in primary UM patients (n =49; Log Rank; p= 0.012) (C) High expression of CysLT_1_ (green) is associated with reduced overall survival in primary UM patients (n =49; Log Rank; p= 0.011) (D) Kaplan-Meier survival curve stratified based by high (green) or low (blue) CysLT_2_ expression and death by metastatic melanoma (n=49; Log Rank; p =0.165). (E) Kaplan-Meier survival curve stratified based by high (green) or low (blue) CysLT_2_ expression and death by any cause (n=50; Log Rank; p =0.332). The third quartile was used as the cut-off point for high versus low expression for all Kaplan-Meier survival curves. Number of events indicates the number of deaths due to metastatic melanoma (B,D). Number of events indicates the number of deaths due to any cause (C,E).

### *2*.*2*. *Cysteinyl leukotriene receptors are expressed in primary and metastatic human uveal melanoma cell lines*

To use UM cell lines as *in vitro* models to investigate the anti-cancer potential of drugs modulating CysLT signalling, we analysed the endogenous expression levels of CysLT_1_ and CysLT_2_ receptors between the primary versus metastatic UM cell lines (Figure 4). Mel285 and Mel270 are derived from primary choroidal melanomas and OMM2.5 is derived from a liver metastasis of the same patient as Mel270 [41]. Both Mel270 and OMM2.5 are GNAQ Q209P positive cell lines, while Mel285 is negative for both GNAQ and GNA11 mutations (Figure 4E) [41]. Real-time PCR confirmed that transcripts for CysLT_1_ and CysLT_2_ are abundantly expressed in all three cell lines with no significant differences in expression levels detected (Figure 4A & B). By Western blotting, prominent 38.5 and 39.6 kDa bands were detected for CysLT_1_ and CysLT_2_, respectively in each cell line (Figure 4C & D). These data confirmed that Mel270, Mel285 and OMM2.5 cell lines were appropriate to analyse the effects of CysLT receptor antagonists on UM cancer hallmarks *in vitro*.

**Figure 4.**
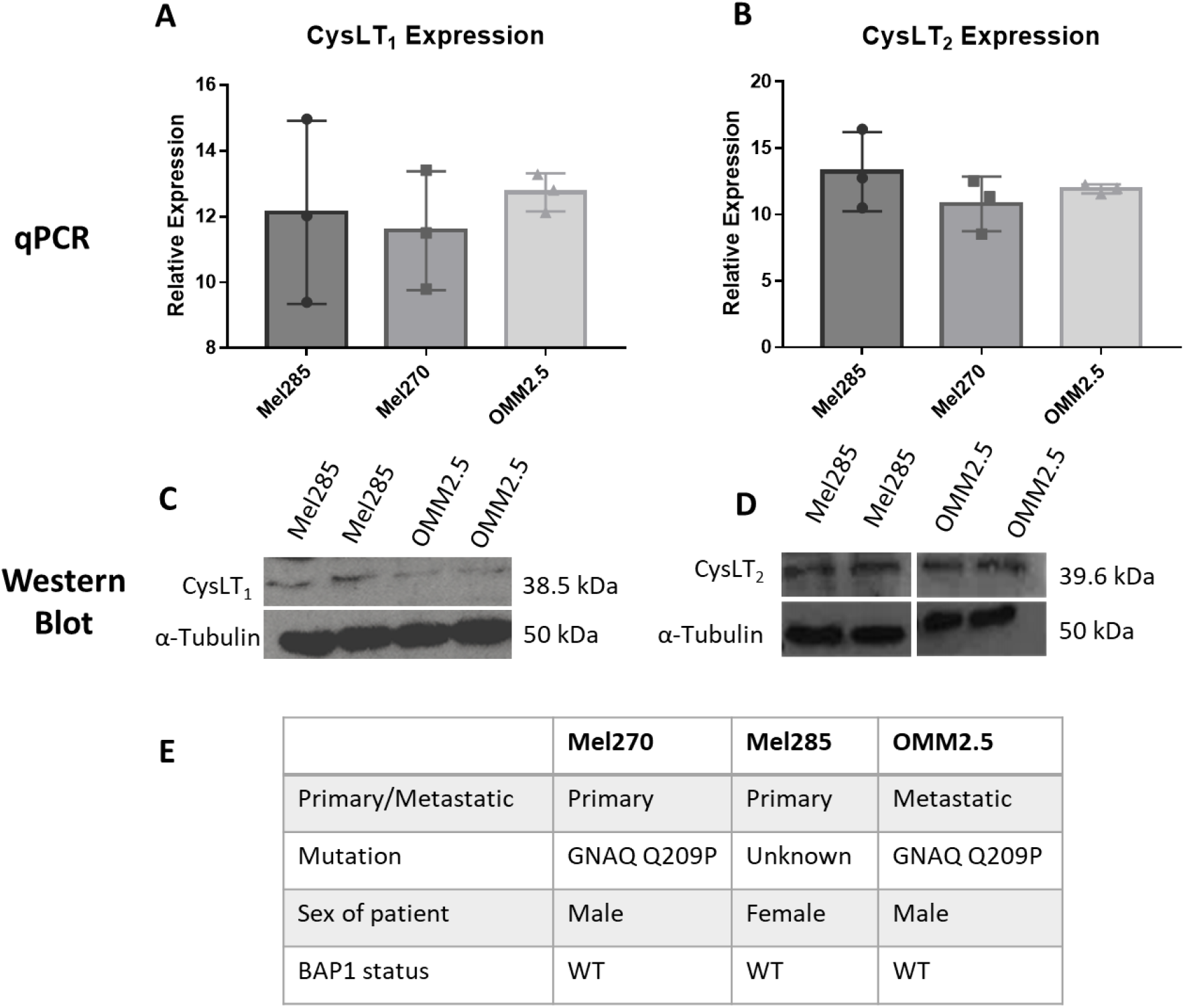
CysLT_1_ and CysLT_2_ are expressed in uveal melanoma cell lines.

qPCR analysis confirmed the expression of CysLT_1_ (A) and CysLT_2_ (B) mRNA in UM cell lines, Mel285, Mel270 and OMM2.5 (n=3). Western blot analysis confirmed the expression of CysLT_1_ (C) and CysLT_2_ (D) in Mel285 and OMM2.5 cells (n=2). (E) Characteristics of UM cell lines used in this study. Statistical analysis was carried out using a paired t-test. Data are expressed as mean + SEM.

### 2.3. CysLT_1,_ but not CysLT_2_, targeting drugs reduce UM cell number in a time- and dose-dependent manner

CysLT receptors activate downstream pathways associated with cell survival and proliferation [42]. Thus, we investigated if CysLT receptor antagonists altered the proliferation of human UM cell lines *in vitro*. Mel270, Mel285 and OMM2.5 UM cell lines were treated with the CysLT_1_-selective antagonists montelukast [43], quininib [32], 1,4-dihydroxy quininib [35]; the CysLT_2_-selective antagonist, HAMI 3379 [44], or dacarbazine, a chemotherapeutic commonly used in the treatment of metastatic UM [45]. In Mel285 cells, after 24 hours of drug treatment, dacarbazine, montelukast and HAMI 3379 did not significantly reduce UM cell number (Figure 5A). In contrast, quininib and 1,4-dihydroxy quininib resulted in a dose-dependent reduction in Mel285 cell number with 50 µM quininib resulting in an average 70.8% reduced cell number (p = 0.029) and 20 µM 1,4-dihydroxy quininib reducing cell number by 61.8% (p = 0.027,) after 24 hours (Figure 5A). In Mel285 cells, after 96 hours of drug treatment, dacarbazine and HAMI 3379 still did not significantly reduce UM cell number (Figure 5E). In contrast to 24-hour treatment, 50 µM montelukast significantly (p = 0.0001) reduced (84% reduction) UM cell number (Figure 5E). Quininib and 1,4-dihydroxy quininib reproduced a dose-dependent reduction in Mel285 cell number with 50 µM quininib now resulting in 95.1% reduced cell number (p = 0.0001) and 20 µM 1,4-dihydroxy quininib now reducing cell number by 55.9% (p = 0.0016) after 96 hours (Figure 5E).

**Figure 5.**
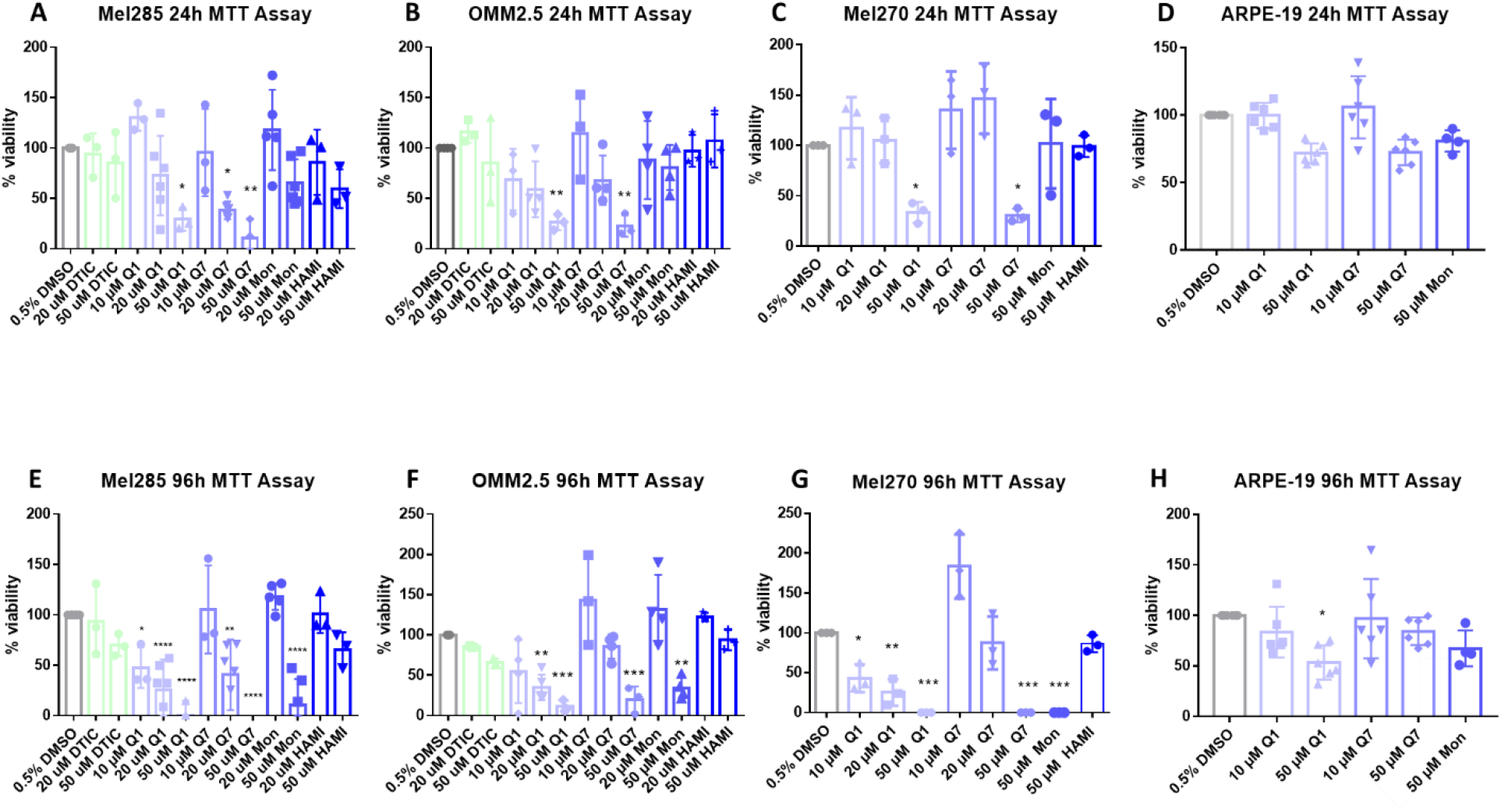
CysLT_1_ antagonists reduce cell viability in uveal melanoma cell lines.

In Mel270 cells, after 24 hours of drug treatment, montelukast and HAMI 3379 did not significantly reduce cell number (Figure 5C). Treatment with quininib and 1,4-dihydroxy quininib resulted in a reduction in Mel270 cell number with 50 µM quininib reducing cell number by 66.4% (p = 0.05) and 50 µM 1,4-dihydroxy quininib reducing cell number by 69.5% (p = 0.0399) (Figure 5C). In Mel270 cells, following 96 hours of drug treatment, HAMI 3379 still failed to significantly reduce cell number (Figure 5G). Treatment with 50 µM montelukast, 50 µM quininib, and 50 µM 1,4-dihydroxy quininib all reduced cell number by 100% (p < 0.0001, for each of the respective treatments) (Figure 5G).

In OMM2.5 cells, a similar trend was observed (Figure 5B & F). Following 24 hours of drug treatment, dacarbazine, montelukast and HAMI 3379 did not significantly reduce cell number (Figure 5B). 50 µM of both quininib and 1,4-dihydroxy quininib significantly (p = 0.0068, p = 0.0043, respectively) reduced (73.5% and 77.1% reductions, respectively) cell viability after 24 hours of treatment. In OMM2.5 cells, after 96 hours of drug treatment, dacarbazine and HAMI 3379 did not significantly reduce cell number. Treatment with 50 µM montelukast reduced cell number by 66% (p = 0.0024). Quininib and 1,4-dihydroxy quininib reproduced a dose-dependent reduction in cell number with 50 µM quininib resulting in an 88.8% reduction (p = 0.0002) and 50 µM 1,4-dihydroxy quininib resulting in a 70.4% reduction (p = 0.0006) (Figure 5F).

To determine whether the effect of CysLT_1_ antagonists was specific to UM cells, ARPE-19 cells, a non-cancerous human retinal pigment epithelium cell line, was treated as a comparator. Following 24 hours of treatment, in contrast to all three UM cell lines where viability was reduced by 60-70%, 50 µM quininib or 1,4-dihydroxy quininib did not significantly reduce (28 and 27.5%, respectively) ARPE-19 cell number (Figure 5A-D). Following 96-hour treatment, 10 µM quininib or 1,4-dihydroxy quininib did not significantly reduce (17 and 3.7%, respectively) ARPE-19 cell number, contrasting with robust effects on viability in UM cells lines (Figure 5D-G). After 96 hours, 50 µM quininib reduced ARPE-19 cell viability by 46.6% (p = 0.0121) (Figure 5H), compared to a 95.1% reduced viability observed in Mel285 cells and an 88.8% reduced viability in OMM2.5 cells. 50 µM 1,4-dihydroxy quininib had no significant effect on ARPE-19 cell number at 96 hours (Figure 5H).

Our results suggest that drugs targeting CysLT_1_, but not CysLT_2_, selectively alters primary and metastatic UM cell number in a time- and dose-dependent manner.

Graphs represent the effects of treatment with varying concentrations of dacarbazine (DTIC), quininib (Q1), 1,4-dihydroxy quininib (Q7), montelukast (Mon) and HAMI 3379 (HAMI) for 24 and 96 hours in Mel285 (A, E) (n=3/n=6), OMM2.5 (B, F), (n=3/n=4), Mel270 (C, G) (n=3) and ARPE-19 (D, H) (n=4/n=6) cell lines. Treatment with 50 μM of quininib, 1,4-dihydroxy quininib, and montelukast for 96 hours significantly reduced cell viability in Mel285 (E), OMM2.5 (F), and Mel270 (G) cells. (D) Quininib analogues had no effect on ARPE-19 cells at 10 or 50 μM following 24 treatment. (H) 50 μM of quininib reduced ARPE-19 viability at 96 hours. Viability of cells was determined using MTT (3-(4,5-dimethylthiazol-2-yl)-2,5-diphenyltetrazolium bromide) assay. 5,000 cells were seeded and treated in triplicate for each individual experiment. Statistical analysis was performed by ANOVA with Dunnett’s post hoc multiple comparison test. Error bars are mean +S.E. *p < 0.05; **p < 0.01; ***p < 0.001; **** p < 0.0001

### 2.4. Quininib drugs inhibit long-term UM cell proliferation

Clonogenic colony formation assays were conducted in Mel285 and OMM2.5 cell lines to determine if CysLT receptor drugs attenuate long-term UM cell proliferation. Mel270 cells failed to grow when seeded at the low cell density required for this assay and as a result, were excluded. UM cells were treated for 24 - 96 hours and grown for 10 additional days prior to analysis of clone survival (Figure 6).

**Figure 6.**
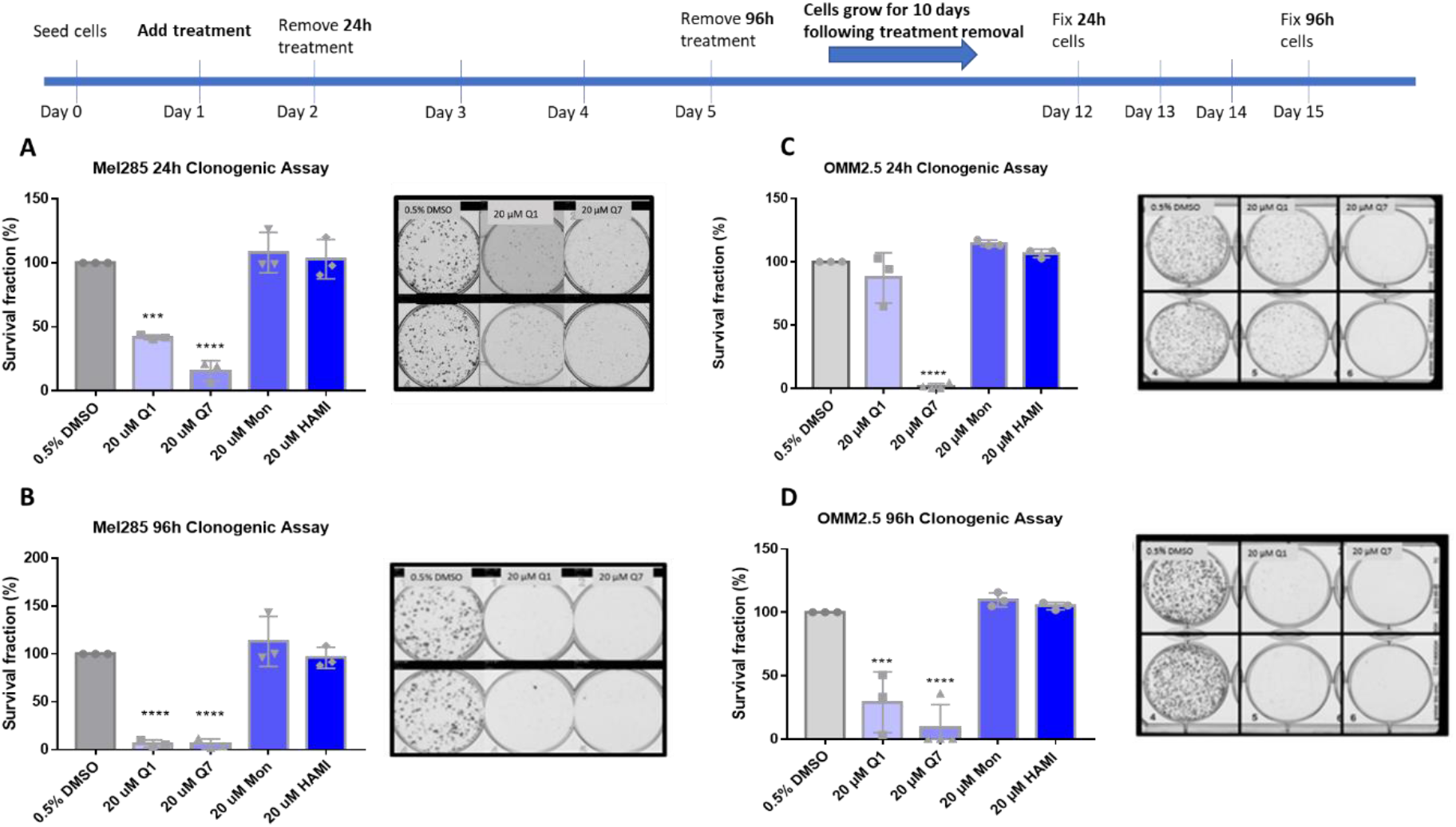
Quininib and 1,4-dihydroxy quininib reduce colony formation in Mel285 and OMM2.5 cells.

In Mel285 cells, UM clone survival after 24 hours treatment was not significantly reduced by treatment with 20 uM montelukast or HAMI 3379. 24-hour treatment with quininib significantly reduced clone survival by 58.1% (p = 0.0002) (Figure 6A). Likewise, 20 uM 1,4-dihroxy quininib significantly reduced Mel285 clone survival by 84.3% (p = 0.0001) after 24 hours treatment. 20 uM montelukast or HAMI 3379 failed to significantly reduce clone survival following treatment for 96 hours. The trend observed with 20 uM quininib and 1,4-dihydroxy quininib was reproduced following 96-hour treatment. 20 uM quininib reduced clone survival by 93.7% (p = 0.0001) and 20 uM 1,4-dihydroxy quininib reduced clone survival by 94% (p = 0.0001) (Figure 6B). There was a statistically significant reduction in clone survival in Mel285 cells treated with quininib for 96 hours compared to those treated for 24 hours (p = 0.0064) (Supplementary Figure 2A).

In the metastatic UM cell line OMM2.5, 20 uM 1,4-dihydroxy quininib was the only drug to significantly reduce clone survival following 24 hours of treatment (98.2% reduction, p = 0.0001) (Figure 6C). Treatment with 20 uM quininib, montelukast or HAMI 3379 for 24 hours did not significantly alter clone survival. In contrast, following 96-hour treatment with quininib a 78.2% reduction (p = 0.0004) was observed. 96-hour treatment with 1,4-dihydroxy quininib produced a 91% reduction (p = 0.0001). Treatment with 20 uM montelukast or HAMI 3379 did not significantly alter clone survival in OMM2.5 cells (Figure 6D). Consistent with the Mel285 data, there was a statistically significant decrease in clone survival in OMM2.5 cells treated with quininib for 96 hours when compared to those treated for 24 hours (p = 0.0164) (Supplementary Figure 2B).

Interestingly, there were some significant differences between cell lines. Following 24 hours treatment, 20 uM quininib reduced clone survival of Mel285 cells by 58.1% compared to a non-significant reduction in OMM2.5 cells (p = 0.0167) (Supplementary Figure 2C). In contrast, treatment with 20 uM 1,4-dihydroxy quininib for 24 hours reduced clone growth by 98.2% in OMM2.5 cells compared to 84.3% in Mel285 cells (p = 0.0180) (Supplementary Figure 2C).

In summary, the CysLT_1_ -selective antagonists quininib and 1,4-dihydroxy quininib are effective at inhibiting long-term proliferation of primary and metastatic UM cell lines. In both cell lines, quininib is more effective following 96 hours of treatment. In Mel285 cells, 20 uM quininib is more effective following 24-hour treatment while in OMM2.5 cells, 20 uM 1,4 -dihydroxy quininib is more effective following 24-hour treatment.

Graphs show the percentage survival fraction of clones at 24 (A, C) and 96 hours (B,D) post treatment. Images of clones captured by GelCount™ system (Oxford Optronix) after 10 days of culture following treatment with DMSO control or 20 μM quininib (Q1) or 20 μM 1,4-dihydroxy quininib (Q7) for 24 or 96 hours. Clones were stained with 0.5% crystal violet before counting. 1,500 cells (Mel285) or 9,000 cells (OMM2.5) were seeded and treated in duplicate in 6-well plates for each individual experiment and individual experiments were conducted three times (n = 3). Statistical analysis was performed by ANOVA with Dunnett’s post hoc multiple comparison test. Error bars are mean +S.E. *p < 0.05; ***p < 0.001; **** p < 0.0001.

### 2.5. Quininib drugs alter the cancer secretome of inflammatory and angiogenic factors in Mel285 and OMM2.5 UM cell lines

We hypothesised that the effects of CysLT receptor drugs on UM cell proliferation and survival may be mediated via modulating angiogenic or inflammatory pathways. Multiplex ELISA quantified whether the secretion of a panel of inflammatory (IFN-γ, IL-10, IL-12p70, IL-13, IL-1β, IL-2, IL-4, IL-6, IL-8, TNF-α) (Figure 7) and angiogenic markers (bFGF, Flt-1, PIGF, Tie-2, VEGF-C, VEGF-D and VEGF-A) (Figure 8) was altered in conditioned media from Mel285 and OMM2.5 cells treated for 24 hours. For both cell lines, secreted PIGF, TIE-2 and VEGF-D were under the level of detection and were removed from analysis (data not shown). IFN-γ and IL-4 secretion were not significantly altered in either cell line following treatment with any tested drugs (Supplementary figure 3).

**Figure 7.**
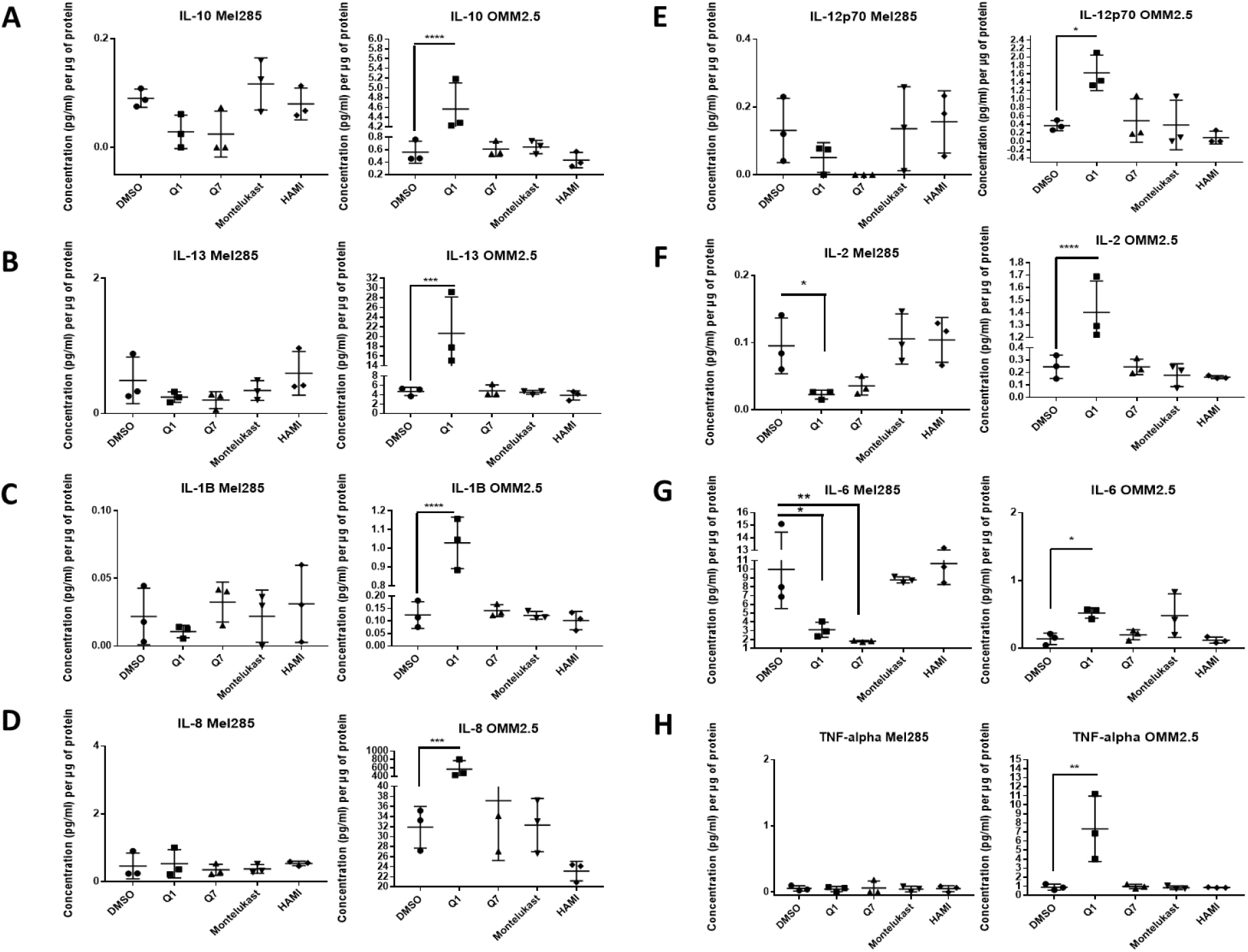
ELISA of cell conditioned media demonstrates that 24-hour treatment with 20 μM quininib analogues decreases inflammatory markers in Mel285 cells but increases inflammatory markers in OMM2.5 cells.

**Figure 8.**
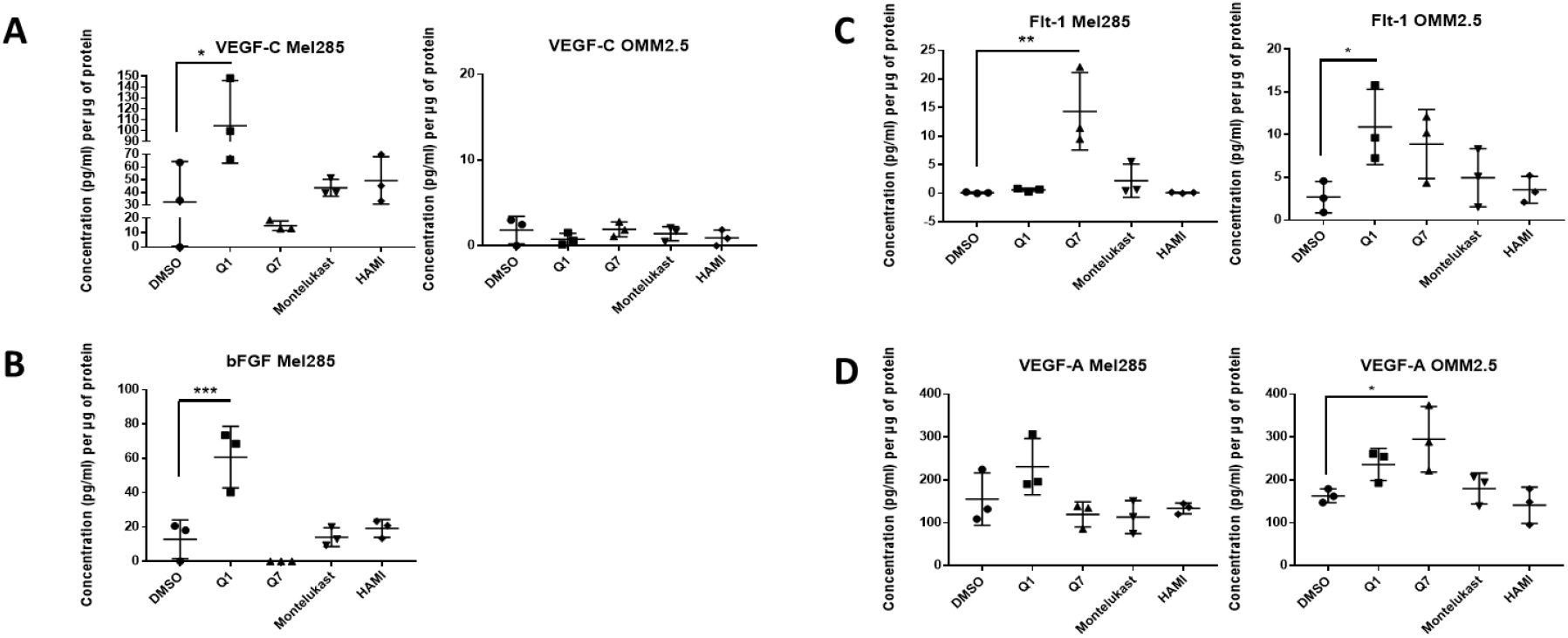
ELISA of cell conditioned media demonstrates that 24-hour treatment with 20 μM quininib analogues increases angiogenic markers in Mel285 cells and OMM2.5 cells.

However, in Mel285 cells, treatment with 20 uM quininib significantly reduced secreted levels of IL-2 and IL-6 (p = 0.0437, p = 0.0142, respectively), and significantly increased secreted levels of bFGF and VEGF-C (p = 0.0006, p = 0.0179 respectively) (Figure 7 & 8). Treatment with 20 uM 1,4-dihydroxy quininib in Mel285 cells significantly decreased secreted IL-6 (p = 0.0048), and significantly increased secreted Flt-1 (p = 0.0014) (Figure 7 & 8). Treatment with 20 uM montelukast or 20 uM HAMI 3379 showed no statistical difference on the secretion of any inflammatory or angiogenic markers examined in Mel285 UM cells (Figure 7 & 8, Supplementary Figure 3).

In the metastatic OMM2.5 cell line, treatment with 20 uM quininib significantly upregulated the secretion of IL-10, IL-1β, IL-2 (p = 0.0001), IL-13 (p = 0.0007), IL-8 (p = 0.001), IL-12p70 (p = 0.0119), TNF-α (p = 0.0023), IL-6 (p = 0.0441)and Flt-1 (p = 0.0362) (Figure 7). Treatment with 20 uM 1,4-dihydroxy quininib significantly upregulated the secretion of VEGF-A (p = 0.0183) (Figure 8). Again, treatment with 20 uM montelukast or 20 uM HAMI 3379 showed no statistical difference on the secretion of any inflammatory or angiogenic markers examined in OMM2.5 cells (Figure 7 & 8, Supplementary Figure 3).

The quininib drugs also produce cell line-dependent effects on the cancer secretome of UM cell lines. For example, in Mel285 cells, quininib decreases the secretion of inflammatory factors (IL-2 and IL-6) and increases the secretion of angiogenic factors (bFGF and VEGF-C). In OMM2.5 cells, a similar upregulation of angiogenic factors was observed following treatment with quininib (Flt-1 and VEGF-A) but in profound contrast, quininib increased the secretion of 8 inflammatory factors (IL-10, IL-12p70, IL-13, IL-1β, IL-2, IL-6, IL-8, TNF-α) in OMM2.5 cells.

In OMM2.5 cells, 24-hour treatment with quininib (Q1) significantly increased the secretion of IL-10 (A), IL-13 (B), IL-1B (C), IL-8 (D), IL-12p70 (E), IL-2 (F), IL-6 (G), and TNF-alpha (H). In Mel285 cells, 24-hour treatment with 20 μM quininib (Q1) and 1,4 – dihydroxy quininib (Q7) significantly reduced the levels of IL-2 (F) and IL-6 (G). Treatment with 20 μM montelukast or HAMI 3379 had no effect on the secretion of inflammatory markers in either cell line. Conditioned media was collected from three separate experiments and analysed by ELISA (n=3). All secretions were normalised to total protein content. Statistical analysis was performed by ANOVA with Dunnett’s post hoc multiple comparison test. Error bars are mean +S.E. *p < 0.05; ** p < 0.01; ***p < 0.001; **** p < 0.0001.

In Mel285 cells, 24-hour treatment with 20 μM quininib (Q1) significantly increased the secretion of VEGF-C (A) and bFGF (B). Treatment with 20 μM 1,4-dihydroxy quininib (Q7) significantly increased the secretion of Flt-1 (C). In OMM2.5 cells, 24-hour treatment with quininib significantly increased the secretion of Flt-1 (C) and VEGF-A (D). Treatment with 20 μM montelukast or HAMI 3379 had no effect on the secretion of inflammatory markers in either cell line. Conditioned media was collected from three separate experiments and analysed by ELISA (n=3). All secretions were normalised to total protein content. Statistical analysis was performed by ANOVA with Dunnett’s post hoc multiple comparison test. Error bars are mean +S.E. *p < 0.05; ** p < 0.01.

### 2.6. CysLT_1_ antagonists inhibit oxidative phosphorylation, but not glycolysis, in UM cell lines

Altered metabolism is a hallmark of cancer [46] and oxidative phosphorylation has emerged as a therapeutic strategy [47]. Comparing 31 tumour types, UM ranked amongst the tumours with the highest oxidative phosphorylation signature [48]. To determine if the effects of CysLT_1_ antagonists were due to altered metabolism in UM cells, the Seahorse Assay was performed assessing two key metabolic pathways, oxidative phosphorylation, and glycolysis. Mel285 and OMM2.5 cell lines were treated for 24 hours with 20 µM of quininib, 1,4-dihyroxy quininib or montelukast prior to quantification of live metabolic readout. Quininib and 1,4-dihyroxy quininib significantly inhibit oxidative phosphorylation in Mel285 cells (p = 0.001 and p = 0.05, respectively) (Figure 9A). In OMM2.5 cells, quininib, 1,4-dihyroxy quininib and montelukast exert significant inhibitory effects on oxidative phosphorylation (p = 0.01, p = 0.05 and p = 0.05 respectively) (Figure 9G). Quininib has a significantly greater effect on the reduction of maximal respiration in OMM2.5 cells compared to Mel285 cells (p < 0.05, data not shown). None of the drugs tested alter glycolysis in either cell line.

**Figure 9.**
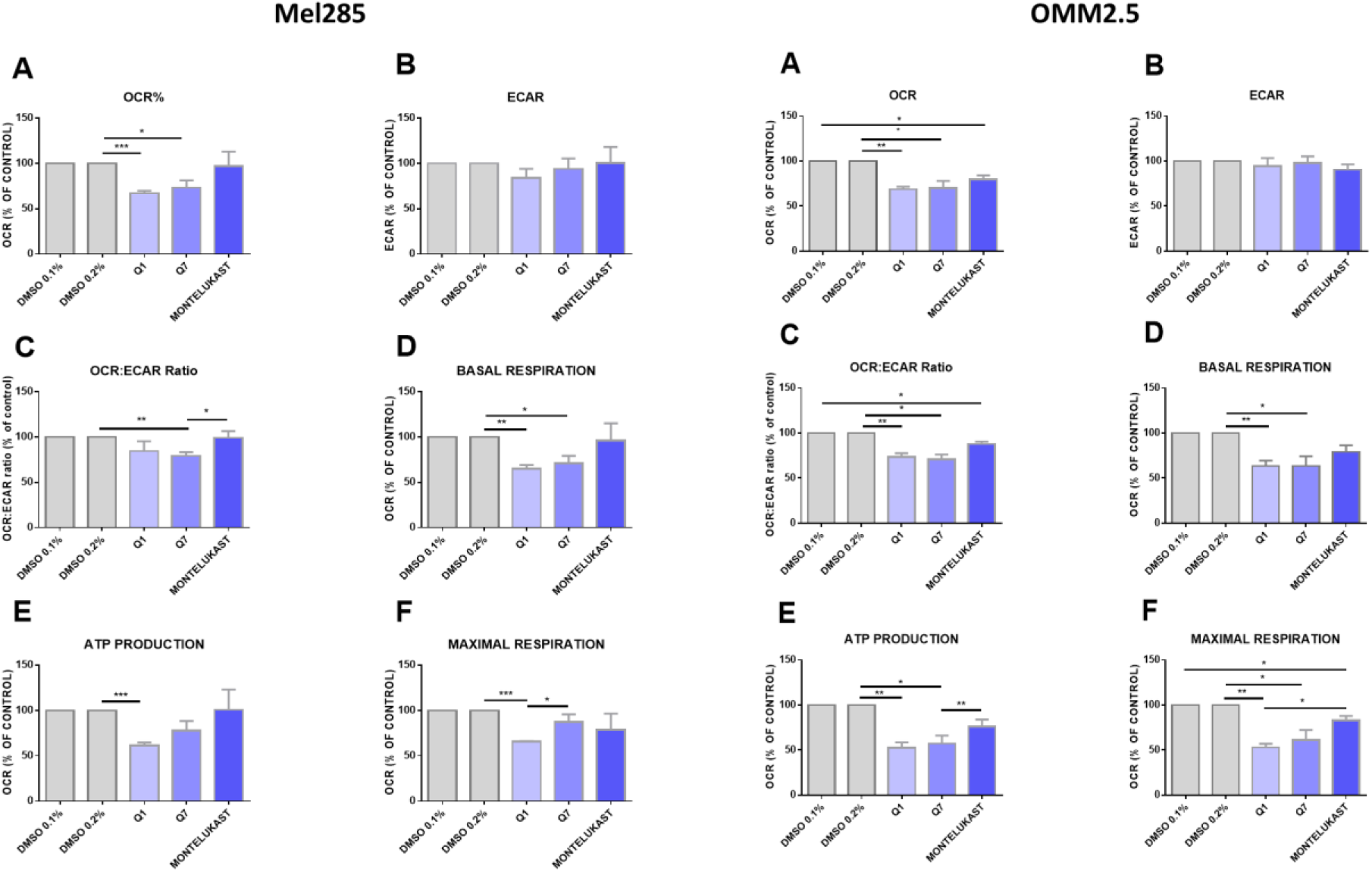
Quininib analogues inhibit oxidative phosphorylation, but not glycolysis, following 24-hour treatment in Mel285 and OMM2.5 cell lines.

Analysis of cellular metabolism in Mel285 (A-F) and OMM2.5 (G-L) UM cell lines. Oxygen consumption rate (OCR), a measure of oxidative phosphorylation, was evaluated in Mel285 (n=5) (A) and OMM2.5 (n=4) (G) cells using the Seahorse Biosciences XFe24 analyser following 24-hour treatment with 20 μM of test compound or DMSO control. (A) 20 μM quininib (Q1) and 20 μM 1,4-dihydroxy quininib (Q7) significantly reduced OCR in Mel285 cells versus DMSO control. (G) 20 μM quininib, 1,4-dihydroxy quininib, and montelukast significantly reduced OCR in OMM2.5 cells versus DMSO control. Extracellular acidification rate (ECAR), a measure of glycolysis was evaluated in Mel285 (B) and OMM2.5 cells (H) following 24-hour treatment. OCR:ECAR ratio was measured in Mel285 (C) and OMM2.5 (D) cells following 24-hour treatment. Basal respiration was significantly reduced in Mel285 (D) and OMM2.5 (J) cells following 24-hour treatment with quininib analogues at 20 μM. (E) ATP production was significantly reduced in Mel285 cells following 24-hour treatment with quininib. (K) ATP production was significantly decreased in OMM2.5 cells following 24-hour treatment with quininib and 1,4-dihydroxy quininib. (F) Maximal respiration was significantly reduced in Mel285 cells following 24-hour treatment with quininib. (L) Maximal respiration was significantly reduced in OMM2.5 following 24-hour treatment with quininib, 1,4-dihydroxy quininib, and montelukast. Data are expressed as mean + SEM. Statistical analysis was carried out using a paired t-test to compare within the same cell line. Data was normalised to cell number, as assessed by crystal violet assay. *p < 0.05; **p < 0.01; ***p < 0.001.

### 2.7. CysLT_1_ antagonists inhibit the in vivo growth of UM xenografts in zebrafish

To determine if the *in vitro* attenuation of UM cell proliferation with CysLT_1_ antagonists could be reproduced *in vivo*, we sought to generate mouse and zebrafish UM cell xenografts [49]. In our experience, Mel285 cell lines were not amenable to generate murine xenograft models following subcutaneous, intraocular, intrahepatic, or intravenous implantation (data not shown). In contrast, the metastatic OMM2.5 cell line produced tumours following subcutaneous, intraocular, or intrahepatic implantation. However, this required lengthy growth times of 7-8 months for the initial subcutaneous cell suspension injection, and 3-4 months for propagation from corresponding subcutaneous fragment implants (Supplementary Figure 4). OMM2.5 cells implanted intraocularly give rise to ocular tumours 3-4 months post cell suspension injection (Supplementary Figure 4A & B). Similarly, OMM2.5 cells implanted intrahepatically allow tumour growth 3-4 months after cell suspension injection (Supplementary Figure 4C & D), and 1-2 months after the re-implantation of tumour fragments arising (Supplementary Figure 4E). Histological examination of the tumours confirmed the presence of a UM cell phenotype (Supplementary Figure 4 F,G & H). Further characterisation of these murine UM xenografts models will ascertain their value in researching the therapeutic effect of UM treatments.

The zebrafish xenograft models proved to be more time-and cost-effective and thus allowed investigation of the CysLT_1_ antagonists on UM cell lines *in vivo*. Mel285 and OMM2.5 UM cell lines were injected into the perivitelline space or eye of 2 dpf zebrafish larvae. Following injection, the larvae were treated with the maximum tolerated dose of quininib (3 µM), 1,4-dihyroxy quininib (10 µM) or montelukast (20 µM). Montelukast significantly reduced the tumour size of Mel285 xenografts propagated in the perivitelline space (p<0.0498, 14.6% reduction) or eye (p<0.0001, 32.4% reduction) (Figure 10A). In contrast, montelukast had negligible effect on OMM2.5 xenografts in the perivitelline space but modestly reduced (p<0.0439 and 25.6% reduction) tumour size in the eye (Figure 10B). In relation to quininib and 1,4-dihydroxy quininib, the most significant reductions in tumour size were also observed with the Mel285 cells line xenografts in the eye (p<0.0001 and 26.2% reduction versus p<0.0001 and 21.7% reduction, respectively) (Figure 10A). Quininib and 1,4-dihydroxy quininib had negligible effects on OMM2.5 xenografts into the perivitelline space, whereas 1,4-dihydroxy quininib modestly reduced (p<0.0258 and 18.4% reduction) OMM2.5 xenograft growth in the eye (Figure 10B). In summary, CysLT_1_ antagonists can inhibit the *in vivo* growth of UM cancer cells; however, the effects are more pronounced with the primary Mel285 compared to the metastatic OMM2.5 cell line. Similarly, the drugs seem to have a more pronounced effect on cells implanted into the eye, the physiologically relevant organ, rather than those implanted into the perivitelline space.

**Figure 10.**
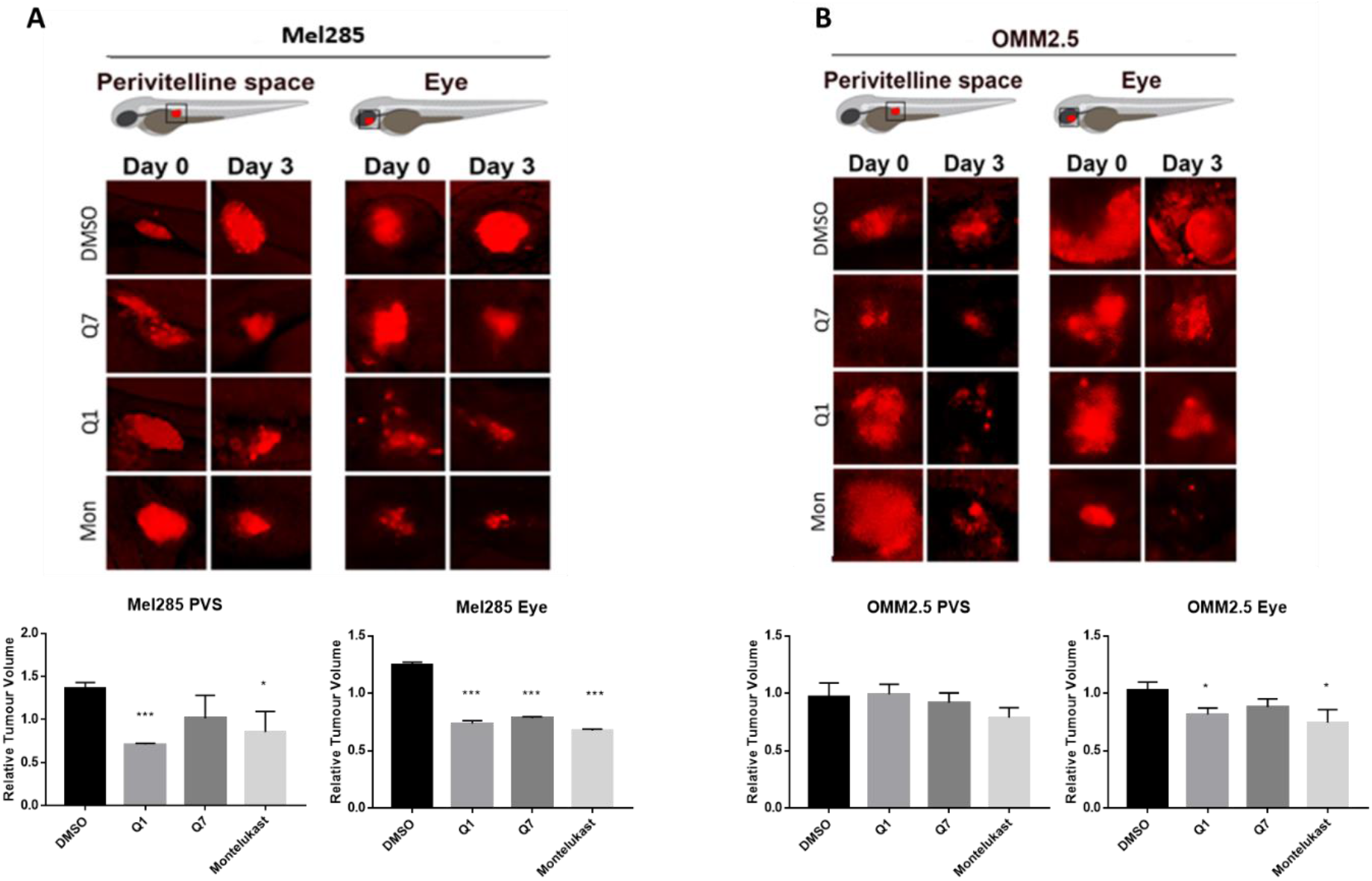
CysLT_1_ antagonists inhibit the growth of UM cell lines in *in vivo* zebrafish xenograft models.

Zebrafish cell line-derived xenograft models were developed using Mel285 and OMM2.5 cell lines. Labelled UM cells were implanted into the perivitelline space (PVS) or vitreous of 48 hpf *Tg(fli1a:EGFP)*^y1^ zebrafish embryos and treated with the maximum tolerated dose of test compound, or DMSO control. (A) Treatment with 3 μM quininib (Q1), 10 μM 1,4-dihydroxy quininib (Q7) and 20 μM montelukast significantly reduced the growth of Mel285 xenografts in the zebrafish eye (p<0.0001). (B) Treatment with 10 μM 1,4-dihydroxy quininib and 20 μM montelukast significantly reduced the growth of OMM2.5 xenografts in the zebrafish eye (p = 0.0258 and p = 0.0439, respectively). Relative change in tumour volume was evaluated as the size of the tumours at three days post implantation (3 dpi/5 dpf) relative to the size immediately after implantation (at 0 dpi/2 dpf).

## 3. Discussion

We identified that high CysLT_1_ expression is associated with survival of primary UM patients and that antagonism of CysLT_1_ alters cancer hallmarks in UM cells *in vitro* and *in vivo*.

To our knowledge, this is the first study to investigate the link between the expression of CysLT receptors in UM patient samples and associated clinical data. Gene expression data from TCGA suggest that high expression of *CYSLTR1* or *CYSLTR2* is significantly linked to disease-free and overall survival in UM patients. This is supported by the known importance of the CysLT_2_/G_αq/11_/PLCB4 pathway in UM oncogenesis. *CYSLTR2* acts as an oncogene [12], albeit in a small subset of UM. Activation of this receptor, and the associated downstream signalling pathways, are identified drivers of disease progression in early UM [50].

TCGA data suggest that high expression of *CYSLTR1* is associated with a poor patient prognosis in UM. This finding was supported by the data generated at the protein level using the patient UM TMA, whereby CysLT_1_ was significantly associated with reduced melanoma-specific survival and reduced overall survival in a primary UM patient cohort. CysLT_1_ was found to be highly expressed in primary UM, with 52/52 tissue samples staining positive for the receptor.

CysLT_2_ was highly expressed in all primary UM examined, with 51/51 tissue samples staining positive for the receptor. All patient samples stained were found to be positive for both CysLT_1_ and CysLT_2_. Both receptors are not highly expressed in normal uveal tract [51], which suggests a role in early carcinogenesis. Of note, high CysLT_1_ expression is associated with poor prognosis and reduced survival in both colorectal [28] and breast cancer [29], while CysLT_2_ has been reported to have an anti-tumorigenic effect in colorectal cancer [52].

Interestingly, high expression of CysLT_1_ was also significantly associated with ciliary body involvement, a feature associated with an increased risk of metastasis [53]. This further suggests a potential link between CysLT_1_ expression and patient prognosis. Given their link to the MAPK pathway, and the known importance of this pathway in UM, it is not surprising that CysLT receptors are highly expressed in primary UM. Similarly, based on the biology of CysLT receptors it is unsurprising that high expression of both CysLT_1_ and CysLT_2_ in UM are associated with significant alterations in inflammation and angiogenesis. Indeed, their high level of expression and link to prognostic and clinical features of the disease may suggest that they are associated with malignant transformation of the tumour. Thus, further investigation using additional UM patient samples is warranted to determine if high CysLT_1_ or CysLT_2_ expression can be statistically associated with patient prognosis and metastatic disease development.

As CysLT receptors are druggable G-protein coupled receptors, we hypothesised that pharmacological antagonists of these receptors may attenuate cancer phenotypes of UM cells in culture. qPCR and western blot analysis confirmed equivalent expression of both receptors in Mel285 and OMM2.5 cell lines suggesting that phenotypic differences observed following drug treatment are not due to differences in CysLT receptor expression. The CysLT_2_ antagonist HAMI 3379 was previously tested in Leu129Gln CysLT_2_ UM oncogene models but showed limited activity as an inverse agonist in these models of constitutive CysLT_2_ activation [54]. To our knowledge, no previous studies have investigated the anti-cancer potential of CysLT_1_ antagonists in UM. However, montelukast exerts anti-cancer activity in chronic myeloid leukaemia [55], colorectal cancer [56], lung cancer [57], and is chemopreventative against 14 different cancer types [23]. Likewise quininib and 1,4 dihydroxy quininib have significant anti-cancer properties in colorectal cancer models [34,35]. Here, we uncover that CysLT_1_ antagonists, but not CysLT_2_ antagonists, produce significant anti-cancer effect in primary and metastatic UM cell lines through the inhibition of cell survival and cell proliferation. The 96 hours of drug treatment in short-term cell viability and long-term cell proliferation assays revealed interesting results. The CysLT_2_ antagonist HAMI 3379 had no effect on the viability of the primary or metastatic UM cell lines. All three CysLT_1_ antagonists significantly reduced viability of the primary and metastatic UM cells, with no intra-drug differences across the cell lines, but with quininib being the most potent in both cell lines. Dacarbazine, a chemotherapeutic clinically used in treatment of metastatic UM, did not reduce the viability of either UM cell line. This is consistent with clinical findings wherein dacarbazine, did not offer any survival advantage in treating metastatic disease [6] or in the adjuvant setting after primary tumour resection [58]. Of note, here, all CysLT_1_ drugs tested performed significantly better than dacarbazine. In long-term proliferation assays the CysLT_2_ antagonist HAMI 3379 again exerted negligible effect on the primary or metastatic UM cell lines. The CysLT_1_ antagonist quininib and 1,4-dihydroxy quininib significantly attenuated proliferation of both cell lines with a seemingly higher efficacy in the primary UM cell line. The same effect of CysLT_1_ antagonists is not observed in ARPE-19 cells, suggesting that this is a specific effect in UM cells.

Here, we identify quininib and 1,4-dihydroxy quininib to significantly alter the secretion of cancer-associated inflammatory and angiogenic factors from UM cell lines. In agreement, the anti-angiogenic and anti-inflammatory actions of CysLT_1_ antagonists are known in other contexts [32,33,59]. CysLTs modulate vascular permeability through upregulation of VEGF expression [60] and CysLT_1_ antagonists modulate vascular permeability through reduction of VEGF expression in mice [61] and in human asthmatic patients [62]. Likewise, the eyes of UM patients often contain increased levels of inflammation-related cytokines in the aqueous humor [63] and several *in vitro* studies show the expression of soluble inflammatory factors in UM. Our TCGA data analysis also suggests that high expression of CysLT receptors is significantly linked to pro-inflammatory and pro-angiogenic pathways. In relation to secreted inflammatory and angiogenic factors, again montelukast and HAMI 3379 had negligible effects. It has been reported that blocking pro-angiogenic isoforms of VEGF, through inhibition of SRPK1, inhibits melanoma tumour growth *in vivo* [64]. Quininib and 1,4-dihydroxy resulted in significant, but dramatically differential effects on the cell lines and factors. For example, quininib treatment was associated with a significant increase in VEGF-C secretion from Mel285 cells, but not from OMM2.5 cells. 1,4-dihydroxy quininib had negligible effects on VEGF-C secretion, but significantly increased Flt-1 in Mel285 in contrast to quininib which had no effect. Interestingly, Flt-1 negatively modulates angiogenesis via its actions as a decoy receptor by trapping VEGF and preventing its binding to VEGFR-2 [65]. Further investigation is required to determine if the drug-induced changes in angiogenic factor expression are cause-or-effect for the corresponding reduction in UM cell viability and proliferation.

Profound differences in the secretion of inflammatory markers was also observed with the quininib drugs. For example, quininib significantly reduced the secretion of IL-2 in Mel285 cells, but significantly increased IL-2 secretion from OMM2.5 cells, whereas 1,4 -dihydroxy quininib exerted negligible effects on IL-2 in OMM2.5 cells. Particularly noticeable is the significantly increased secretion of 7 inflammatory factors in OMM2.5, but not Mel285, after quininib, but not 1,4-dihydroxy quininib, treatment. Notably, 24 hours treatment with quininib reduced the survival and proliferation of Mel285 but not OMM2.5 cells. This is potentially due to a resistance associated up-regulation of inflammatory factors in OMM2.5 cells. Indeed, the basal expression of IL-13 and IL-8 is much higher in OMM2.5 than Mel285 cells. The pattern of IL-2 and IL-6 secretion may explain the similar actions of quininib drugs on proliferation in the Mel285 cells. UM cell lines express IL-2R, and production of the IL-2 ligand by tumour infiltrating lymphocytes and macrophages stimulates tumour cell proliferation [66]. IL-6 stimulates tumour cell proliferation and survival through the inhibition of apoptosis and interference with IL-6R signalling leads to decreased UM cell viability [67]. Quininib and 1,4 dihydroxy quininib result in 3-5-fold reductions in secreted levels of IL-2 and IL-6. Changes in IL-2 and IL-6 are not sufficient to explain effects on the OMM2.5 cells, but anti-proliferative effects in these cells may be confounded by the up-regulation of multiple other inflammatory factors.

The importance of dysregulated metabolism in the initiation and progression of cancer is well understood. Although much is known about metabolic rewiring in cutaneous melanoma, few studies focus solely on the metabolic underpinnings of UM. Oxidative phosphorylation is upregulated in invasive melanoma [68] and the metabolic switch of some melanomas to oxidative phosphorylation has been linked to resistance to inhibitors of the MAPK pathway [69]. While CysLT are predominantly known for their role in inflammation and angiogenesis, they are linked to alterations in respiratory activity. LTD_4_ increases mitochondrial metabolic activity and mitochondrial gene transcription in human intestinal epithelial cells and colorectal cancer cells [70]. Significantly decreased oxidative phosphorylation was observed in Mel285 and OMM2.5 cells with quininib or 1,4-dihydroxy quininib and OMM2.5 cells with montelukast. Modulation of oxidative phosphorylation controls proliferation of tumour cells [71] which may explain the effect of CysLT_1_ antagonists on UM cells.

*In vivo* models are preferable to human cell lines to study the complexity of the tumour microenvironment. Zebrafish xenograft models have been established as robust preclinical models in which experimental drugs can be tested [49]. To this end, we created xenograft models using UM cell lines, to determine if the effects of CysLT_1_ antagonists on UM cell survival and proliferation can be recapitulated in a more complex system. Our *in vitro* data is supported by the results generated in zebrafish models whereby CysLT_1_ antagonist drugs significantly inhibit the growth of both Mel285 and OMM2.5 zebrafish xenograft models. Interestingly, CysLT_1_ antagonists have a greater effect in zebrafish orthoxenograft models, in which the cells are implanted into the corresponding anatomical location. Given that the Mel285 cell line originated in the eye, and the OMM2.5 cell line originated in the liver, this may explain the more substantial effect observed in the Mel285 orthoxenograft model. To further validate our results, we have generated an OMM2.5 cell line-derived orthotopic xenograft mouse model of UM. This and patient-derived UM xenografts are considered the most appropriate pre-clinical models for future studies to evaluate the potential of CysLT_1_ antagonist as therapeutics for UM.

## 4. Materials and Methods

### Cell Culture

UM cell lines derived from primary (Mel285, Mel270) and metastatic (OMM2.5) UM were kindly provided by Dr. Martine Jager (Leiden, The Netherlands) [41]. Cell lines were maintained at 37°C/ 5% CO_2_ in RPMI 1640 Medium (Gibco) supplemented with 10% FBS and 2% Penicillin/Streptomycin. ARPE-19 cells were maintained at 37°C/ 5% CO_2_ in DMEM: F12 supplemented with 10% FBS, 1% Penicillin/Streptomycin and 2.5 mM L-Glutamine.

### Drug preparation

Quininib (Q1), 1,4-dihydroxy quininib (Q7) [32,33], montelukast (Sigma - SML0101), HAMI 3379 (Cayman Chemical – 10580) and dacarbazine (Sigma - D2390) were dissolved in 100% DMSO and stored as (10-50 mM) stock solutions. Working solutions (100 μM) were prepared fresh prior to each experiment in complete cell culture medium as described above. Drugs were made to final test concentrations by adding the required volume of the working solution to cells in complete media. 0.1 and 0.2% DMSO were used as controls in Seahorse Assay experiments. 0.5% DMSO was used as a control for all other drug treatment experiments.

### MTT Assay

3-(4,5-dimethylthiazol-2-yl)-2,5-diphenyltetrazolium bromide) dye determined cytotoxic effects in cell lines. Cells were trypsinised using trypsin-EDTA (0.05%) (ThermoFisher Scientific) and centrifuged at 1,200 rpm for 5 minutes at RT. Cell pellets were re-suspended in complete medium, cells seeded into 96-well plates at 5,000 cells/well. After 24 hours adherence, cell medium was removed and replaced with the desired drug concentration. 0.5% DMSO in RPMI 1640 (UM cells) or DMEM (ARPE-19 cells) was used as vehicle controls. Cells were incubated for 24 and 96 hours with drugs. Drug solution was removed, and the wells washed with PBS before adding 90 μl of serum-free medium and 10 μl MTT (3-(4,5-dimethylthiazol-2-yl)-2,5-diphenyltetrazolium bromide) dye to each well. The plate was covered and incubated for 2.5 hours at 37°C. 100 μl of 100% DMSO was added to each well to dissolve the formazan crystals. Absorbance values at 570 nm were determined using a SpectraMax® M2 microplate reader.

### Colony Formation Assay

1.5 × 10^3^ (Mel285) or 9 × 10^3^ (OMM2.5) cells were seeded per well of a 6-well plate and allowed adhere for 24 hours. Cells were treated with the desired concentration of drug for 24 or 96 hours. 0.5% DMSO was used as a vehicle control. Following treatment, the drug solution was removed, and cells grown in complete medium for 10 days. Clones were fixed with 4% paraformaldehyde for 10 minutes and stained using 0.5% crystal violet (Pro-Lab diagnostics PL700) for 2 hours at RT. Clone counting was performed using the GelCount™ system (Oxford Optronix).

### Statistical analysis for drug treatment experiments

Statistical analysis applied GraphPad Prism 7 software (GraphPad, San Diego, CA). Specific statistical tests used are indicated in figure legends. All data are presented as mean ± standard error of the mean (SEM). For all statistical analysis, differences were considered statistically significant at p < 0.05.

### CysLT_1_ and CysLT_2_ qPCR

Total RNA was extracted from UM cells using the *mir*Vana™ miRNA Isolation Kit (ThermoFisher Scientific) as per manufacturer’s instructions. Briefly, cells were trypsinised and washed by gently resuspending in 1 ml of PBS and pelleting at 1,200 rpm prior to lysis and total RNA isolation. Following isolation, total RNA concentration was quantified at 260 nm (Spectrophotometer ND-2000) and samples stored at −80 °C. cDNA was synthesized with the SuperScript II Reverse Transcriptase system (Invitrogen) or the TaKaRa PrimeScript™ RT Reagent Kit, using random hexamers as per supplier’s instructions.

### CysLT_1_ and CysLT_2_ Western Blot

UM cells were seeded at 250,000 cells per well of a 6-well plate and left adhere for 24 hours. Total protein was extracted from cells. Cells were washed in ice-cold PBS and lysed with RIPA buffer (Sigma) supplemented with 200 mM NaF, 100 mM PMSF, 100 mM sodium orthovanadate, 1X protease inhibitor cocktail (Sigma), 1X phosphatase inhibitor cocktail 2 (Sigma) and 1X phosphatase inhibitor cocktail 3 (Sigma). Following lysis, cells were scraped into an Eppendorf and left on ice for 45 minutes, with vortexing at 15-minute intervals. Cells were centrifuged at 14,000 rpm for 30 minutes at 4°C. After centrifugation, the supernatant was collected and stored at -80°C. Protein concentrations were determined using the BCA protein assay kit (Fisher). 15 µg of protein was prepared in 4X sample buffer and 10X reducing agent and separated by 10% SDS/ PAGE, transferred to PVDF membranes (Millipore), and probed with primary antibodies (CysLT_1_ : Abcam [ab151484], 1:1000, CysLT_2_ : Cayman Chemical [CAY120560] 1:1000, Alpha-Tubulin : Santa Cruz, 1:200). Secondary antibodies were anti-mouse IgG HRP-linked (Cell Signalling [7076S] 1:1000), or anti-rabbit IgG HRP-linked (Cell Signalling [7074S] 1:1000). Signal was detected using enhanced chemiluminescence as per manufacturer’s instructions (Pierce™ ECL Western Blotting Substrate).

### The Cancer Genome Atlas gene expression analysis

Gene expression and clinical data from 80 primary UM included in The Cancer Genome Atlas (hereafter TCGA-UM dataset) were collected from the GDC data portal through the R package ‘TCGAbiolinks’. RNA-seq data was downloaded in Fragments Per Kilobase of exon per million fragments Mapped (FPKM) and then converted to log2 scale.

The association between *CYSLTR1* and *CYSLTR2* gene expression and prognosis were assessed by Cox proportional hazard regression models, adjusted by sex and age. Disease Free Survival (DFS) and Overall Survival (OS) were used as end points. The third quartile was used as an optimal cut-off to divide samples into High and Low expression categories. Survival probabilities were plotted on a Kaplan-Meier curve and a Log-rank test was used to compare the two groups. Survival analysis was performed with R package ‘survminer’. Disease free survival is defined as time to metastatic recurrence. Overall survival is defined as death by any cause.

Gene Set Variation Analysis was performed to calculate enrichment scores in functions and pathways ‘Inflammatory response’, ‘INF-γ’, ‘Glycolysis’, ‘Oxidative Phosphorylation’, ‘TNF-α’, ‘Angiogenesis’ and ‘GPCR signalling’ (R package ‘GSVA’). They were manually selected from the Molecular Signatures Database (MSigDB) which includes gene sets from Hallmarks and Biocarta curated pathways. Samples were divided by the third quartile gene expression values of *CYSLTR1* and *CYSLTR2*. For each score obtained, differences were assessed using a non-parametric Wilcoxon test. Differences were considered statistically significant when p-value < 0.05.

### Ethics

This study conformed to the principles of the Declaration of Helsinki and Good Clinical Practice guidelines. Approval for the study was obtained from the Health Research Authority (NRES REC ref 16/NW/0380), and all patients provided informed consent.

All experiments involving the use of rodents were approved by the Ethical Committee of Animal Experimentation of the Parc Científic de Barcelona (PCB) under the procedure number 9928-P1 approved by the Generalitat de Catalunya.

### Tissue samples

A tissue microarray (TMA) was generated from primary UM samples of 52 consented patients treated at the Liverpool Ocular Oncology Centre, with the primary UM samples being stored within the Liverpool Ocular Oncology Biobank (HTA Licence 12020 and HRA REC 16/NW/0380).

### Immunohistochemistry

IHC for CysLT_1_ (Abcam – ab151484) and CysLT_2_ (Cayman Chemical - CAY120560) was performed on 4-μm FFPE sections arranged on the above mentioned TMA using commercial equipment (Leica Bond RXm System; Leica Microsystems Ltd, Milton Keynes, UK) and a detection kit (Bond Polymer Refine Red Detection Kit; Leica Biosystems, Inc., Buffalo Grove, IL, USA) as previously described [72]. Slides were counterstained with hematoxylin and mounted using DPX mountant (Sigma Aldrich, Corp.). Colorectal cancer tissue served as the positive control (Supplementary Figure 1A); negative control was omission of the primary antibody (Supplementary Figure 1A). Slides were scanned using a slide scanner (Aperio CS2; Leica Biosystems, Inc.) and analysed with imaging software (Aperio Image Scope version 11.2; Leica Biosystems, Inc.). Each core was scored based on intensity (0, 1, 2 or 3) and percentage of tumour cells stained. The final score was calculated using the following equation: (scoring intensity x % of cells stained)/n number of samples [72]. The IHC-stained slides were scored by three independent investigators (SEC, HK, KS). Melanoma-specific survival is defined as death from metastatic melanoma. Overall survival is defined as death by any cause.

### Digital slide scanning and automated image analysis

Slides were scanned with an Aperio AT2 digital slide scanner (Leica Biosystem, Milton Keynes, UK) with a 20× lens, and the quality of the images was checked manually before the application of the digital algorithm. Automated digital image analysis was performed using the Visiopharm Integrator System (Visiopharm, Hoersholm, Denmark). A cytoplasmic algorithm from the ONCOTOPIX module (v4.2.2.0, Visiopharm, Hoersholm, Denmark) was fine-tuned for the interpretation of CysLT_1_ and CysLT_2_ staining. H-score was used as the image analysis output, which was calculated using the following formula: [1 × (% of weakly positive cells) + 2 × (% of moderately strong positive cells) + 3 × (% strong positive cells)], where the H-score of 0–100 was generally categorized as low expression, 101–200 as intermediate expression, and 201–300 as high expression of CysLT_1_ and CysLT_2._

### Statistical analysis for immunohistochemistry

Survival time (years) was calculated from the date of first diagnosis until death, or study closure on 29^th^ May 2019. All analyses were carried out using SPSS Statistics v.24 (IBM).

### Mel285 and OMM2.5 ELISA

Cells were seeded at 150,000 cells per well of a 6-well plate and allowed adhere overnight. Cells were treated with 20 μM of quininib, 1-4-dihydroxy quininib, montelukast, HAMI 3379, or 0.5% DMSO as control. All treatments were conducted in duplicate. Following 24 hours treatment, 1 ml of media was removed from each well and stored at –20 °C. Media was processed according to MSD (Meso Scale Discovery) multiplex protocol. To assess angiogenic and inflammatory secretions from cell conditioned media, a 17-plex ELISA kit separated across 3 plates was used (Meso Scale Diagnostics, USA). The multiplex ELISA determined the secreted levels of; IFN-γ, IL-10, IL-12p70, IL-13, IL-1β, IL-2, IL-4, IL-6, IL-8, TNF-α, bFGF, Flt-1, PIGF, Tie-2, VEGF-C, VEGF-D and VEGF-A in cell conditioned media. Assays were run as per manufacturer’s recommendation; an overnight supernatant incubation protocol was used for the Pro-inflammatory Panel 1 with the Angiogenesis Panel 1 assay being run on the same day protocol. Cell conditioned media was run undiluted on all assays as per previous optimisation experiments. Secretion data for all factors was normalised to cell lysate protein content (extracted as described above) by using a BCA assay (Fisher).

### Zebrafish Breeding and Maintenance

All experiments carried out on animals were granted ethical approval by Linköping Animal Research Ethics Committee. Zebrafish were maintained in a 14-h light, 10-h dark cycle in a recirculating water system at 28 °C. Larvae were produced through natural spawning and maintained as previously described [73].

### Zebrafish Cell Line Xenografts

Implantation of Mel285 or OMM2.5 cells into zebrafish embryos followed published protocols [74]. Briefly, cells were labelled for 30 min at 37°C in 6 mg/mL DiI (Sigma) in PBS followed by washing 3x in PBS. Labelled cells were implanted in the perivitelline space or vitreous of 48 hpf *Tg(fli1a:EGFP)*^*y1*^ zebrafish embryos, maintained from the 8-cell stage in 0.003% PTU-containing E3-water (5 mM NaCl, 0.17 mM KCl, 0.33 mM CaCl_2_, 0.33 mM MgSO_4_). Approximately 200-500 cells in 2-5 nL were implanted in each embryo (20 embryos per group). Embryos were transferred to individual wells of 24-well plates containing 0.5 mL quininib, 1-4-dihydroxy quininib or montelukast at 10 mM final concentration in E3-PTU water. Tumour-bearing embryos were imaged using a fluorescent microscope (SMZ1500, Nikon) soon after implantation. Embryos with tumour cells erroneously implanted in the yolk, brain or circulation were removed. Embryos bearing tumours in the perivitelline space or the vitreous were incubated at 36°C for three days and re-imaged. Relative change in tumour volume was evaluated as the size of the tumours at three days post implantation (3 dpi) relative to the size immediately after implantation (at 0 dpi).

### Seahorse metabolism measurements

Mel285 and OMM2.5 were seeded in four wells per treatment group at a density of 12,000 and 14,000 cells per well respectively in a 24-well cell culture XFe24 microplate (Agilent Technologies, Santa Clara, CA, USA) at a volume of 100 μl RPMI and allowed adhere at 37°C and 5% CO_2_ for 5 hours, then an additional 150 μl RPMI added. Twenty-four hours following seeding, cells were treated with 20 μM of quininib, 1-4-dihydroxy quininib or montelukast along with 0.1% and 0.2% DMSO controls. Twenty-four hours following drug treatment, cells were washed with unbuffered DMEM supplemented with 10 mM glucose, 10 mM sodium pyruvate (pH 7.4) and incubated for 1 hour at 37°C in a CO_2_-free incubator. The oxygen consumption rate (OCR) and extracellular acidification rate (ECAR) were measured using a Seahorse Biosciences XFe24 Extracellular Flux Analyser (Agilent Technologies, Santa Clara, CA, USA). Three basal measurements of OCR and ECAR were taken over 24 min consisting of three repeats of mix (three min)/wait (2 min)/measurement (3 min) to establish OCR measurement. Three additional measurements were obtained following the injection of three mitochondrial inhibitors including oligomycin (2 μg/ml) (Sigma Aldrich, Missouri, USA), an uncoupling agent carbonyl cyanide 4-(trifluoromethoxy) phenylhydrazone (FCCP) (5 μM) (Sigma Aldrich, Missouri, USA). and antimycin-A (2 μM) (Sigma Aldrich, Missouri, USA) and ATP turnover was calculated by subtracting the OCR post oligomycin injection from baseline OCR prior to oligomycin addition. Proton leak was calculated by subtracting OCR post antimycin-A addition from OCR post oligomycin addition. Maximal respiration was calculated by subtracting OCR post antimycin addition from OCR post FCCP addition. Non-mitochondrial respiration was determined as the OCR value post antimycin-A addition. All measurements were normalised to cell number using the crystal violet assay, transferring the eluted stain to a 96-well plate before reading.

### Crystal violet assay

Cells were fixed in a 1% glutaraldehyde solution for 15 minutes at room temperature followed by two washes in 100 μl of PBS. Cells were stained with 0.1% crystal violet for 30 minutes at room temperature. Crystal violet was removed by washing twice in water. Plates were allowed airdry overnight, then crystal violet stain was eluted using 1% Triton X-100 solution on a plate shaker for 1 hour. The eluted stain was transferred to a 96-well plate and absorbance read at 595 nm on a Versamax plate reader.

## 5. Conclusions

There is an overwhelming, unmet clinical need for targeted therapies for the treatment of UM. The cysteinyl leukotrienes are established as regulators of inflammation and have recently emerged as novel regulators of angiogenesis, two key processes in UM. For the first time, we examine the clinical relevance of CysLT_1_ and CysLT_2_ expression and in primary UM samples and show a link between high CysLT_1_ expression and primary UM patient survival. Our data highlight the involvement of both receptors with clinical features of their disease and reinforces their link to inflammatory and angiogenic pathways. We determined that antagonist drugs of CysLT_1_, but not CysLT_2_, inhibit the survival and proliferation of primary and metastatic UM cells in a specific, time- and dose-dependent manner. This effect is recapitulated in *in vivo* zebrafish cell line xenograft models. Antagonism of CysLT_1_ in UM cells leads to alterations in the secretion of pro-inflammatory and pro-angiogenic secretions and a decrease in oxidative phosphorylation. The importance of the cysteinyl leukotriene receptor signalling pathway, and antagonism of CysLT_1_, in UM should be further explored.

## Supplementary Materials

The following are available, Figure S1: Patient TMA control tissue. Core-core correlations of manual versus digital pathology analysis. Clinical characteristics of patients included in the TMA, Figure S2: Cell line comparison data for the clonogenic assay, Figure S3: Factors unchanged following treatment in Mel285 and OMM2.5 cells, Figure S4: Generation of OMM2.5 cell line-derived orthotopic xenograft models of UM.

## Author Contributions

Conceptualization, K.S, J. O’S and B.N.K; methodology, K.S, H.K, S.E.C, A.R, W.M.G, J.M.P, J.O’S and B.N.K; validation, K.S, A.B.H, R.SP, S.GM, L.D.J, H.K, S.E.C, A.R, W.M.G; formal analysis, K.S and B.N.K; investigation, K.S, A.B.H, R.SP, S.GM, L.D.J, F.O’C, M.H, H.K, S.E.C, A.R, A.V, R.B; resources, B.N.K, S.E.C, W.M.G, A.V; data curation, K.S; writing—original draft preparation, K.S; writing—review and editing, K.S, J.O’S, S.E.C, J.M.P B.N.K; supervision, B.N.K; project administration, K.S and B.N.K; funding acquisition, K.S and B.N.K. K.S, S.E.C, A.R, W.M.G, J.M.P, J.O’S and B.N.K interpreted the results and provided significant intellectual input. All authors have read and agreed to the published version of the manuscript.

## Acknowledgments

We thank Natalie Coplin for assistance with immunohistochemical staining and Dr. Claudia Aura Gonzalez and Dr. Nebras Al Attar for automated image analysis. We thank Dr. Husvinee Sundaramurthi for proof-reading the manuscript.

