## Supplementary Figures 1-4 for "High Cysteinyl Leukotriene Receptor 1 Expression Correlates with Poor Survival of Uveal Melanoma Patients and Cognate Antagonist Drugs Modulate the Growth, Cancer Secretome, and Metabolism of Uveal Melanoma Cells"

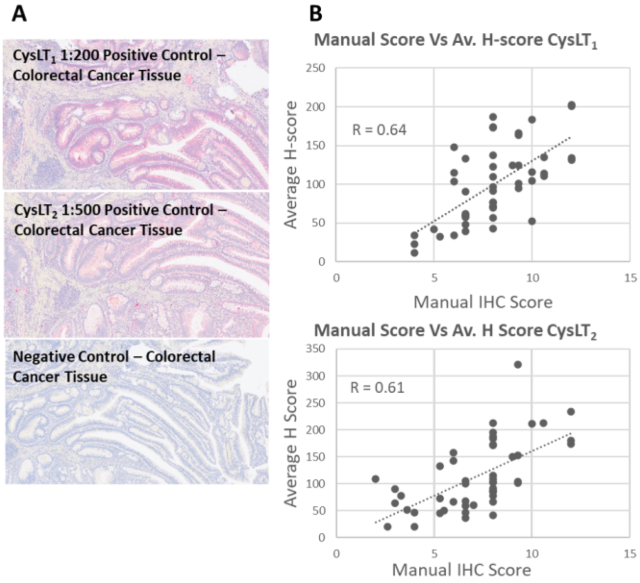


C

| | Age at primary management | Largest ultrasound diameter | Ultrasound height | Mitotic count | Survival (Years) or to study end |
|---|---|---|---|---|---|
| Mean | 62.2 | 17.03 | 9.2 | 6.7 | 8.49 |
| Median | 62.3 | 17.35 | 8 | 4 | 8 |
| Range | 50 | 13 | 13 | 31 | 18.6 |
| Maximum | 39 | 10.6 | 5 | 0 | 0.4 |
| Minimum | 89 | 23.6 | 18 | 31 | 19 |

D

| Clinical Feature | Absent | Present | Unknown | Total number of patients | Statistical correlation with $CysLT_1$ expression | Statistical correlation with $CysLT_2$ expression |
|---|---|---|---|---|---|---|
| Extraocular extension | 49 | 3 | 0 | 52 | 0.963 | 0.11 |
| Monosomy 3 | 28 | 23 | 1 | 52 | 0.187 | 0.889 |
| PAS positive loops | 15 | 37 | 0 | 52 | 0.453 | 0.887 |
| Ciliary body involvement | 37 | 15 | 0 | 52 | **0.041*** | 0.285 |
| Epithelioid cell involvement | 22 | 30 | 0 | 52 | 0.424 | 0.392 |

* $p < 0.05$

***Supplementary Figure 1* Patient TMA control tissue. Core-core correlations of manual versus digital pathology analysis. Clinical characteristics of patients included in the TMA.**

(A) Colorectal cancer tissue positive control for $CysLT_1$. Colorectal cancer tissue positive control for $CysLT_2$. Negative control (omission of antibody). (B) Correlation between manual score and the average H-score assigned by digital pathology analysis for $CysLT_1$ (R=0.64) and $CysLT_2$ (R=0.61). (C) Clinical characteristics of the patients included in the TMA analysed. (D) The relationship between prognostic clinical characteristics in UM and the expression of high $CysLT_1$ or $CysLT_2$. Ciliary body involvement has a statistically significant relationship with high $CysLT_1$ expression ($p = 0.041$).

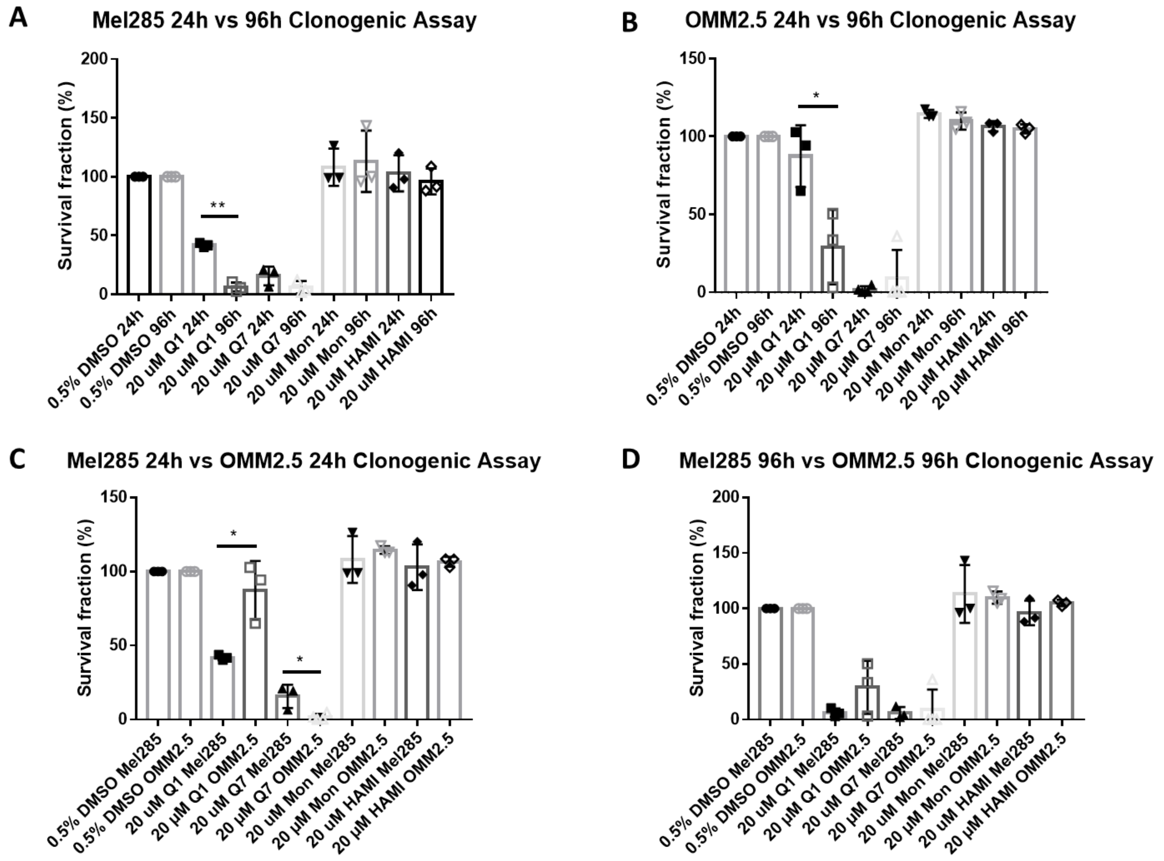


***Supplementary Figure 2* Cell line comparison data for the clonogenic assay**

(A) Differences in the percentage survival fraction of Mel285 clones when treated for 24 versus 96 hours. Treatment with 20 µM quininib (Q1) is significantly more effective at 96 hours. (B) Differences in the percentage survival fraction of OMM2.5 clones when treated for 24 versus 96 hours. Treatment with 20 µM quininib (Q1) is significantly more effective at 96 hours. (C) Differences in the percentage survival fraction of Mel285 versus OMM2.5 clones when treated for 24 hours. 20 µM quininib (Q1) is significantly more effective in Mel285 cells following 24-hour treatment. While 20 µM 1,4 – dihydroxy quininib (Q7) is significantly more effective in OMM2.5 cells following 24-hour treatment. (D) Differences in the percentage survival fraction of Mel285 versus OMM2.5 clones when treated for 96 hours. There were no statistically significant differences between cell lines following 96-hour treatment. Statistical analysis was performed using a paired t-test to compare within the same cell lines. An unpaired t-test was used to compare between different cell lines. Error bars are mean +S.E. $^{*}p < 0.05$; $^{**}p < 0.01$.

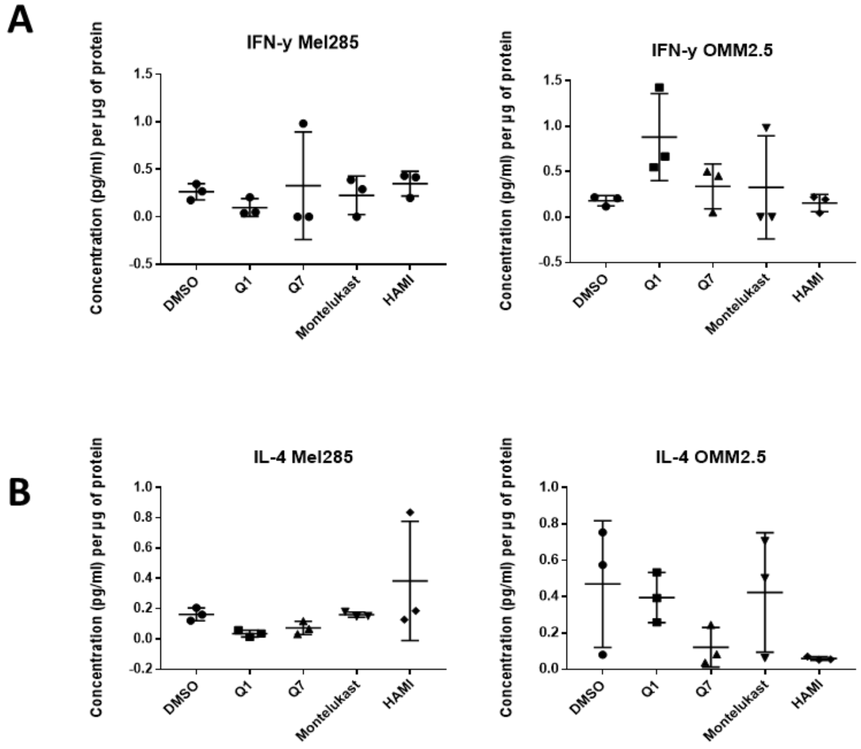


***Supplementary Figure 3* Factors unchanged following treatment in Mel285 and OMM2.5 cells.**

Treatment with all CysLT targeting drugs tested (quininib (Q1), 1,4 –dihydroxy quininib (Q7), montelukast and HAMI 3379 (HAMI)) had no effect on the secretion of IFN- γ (A) or IL-4 (B) in Mel285 or OMM2.5 cells.

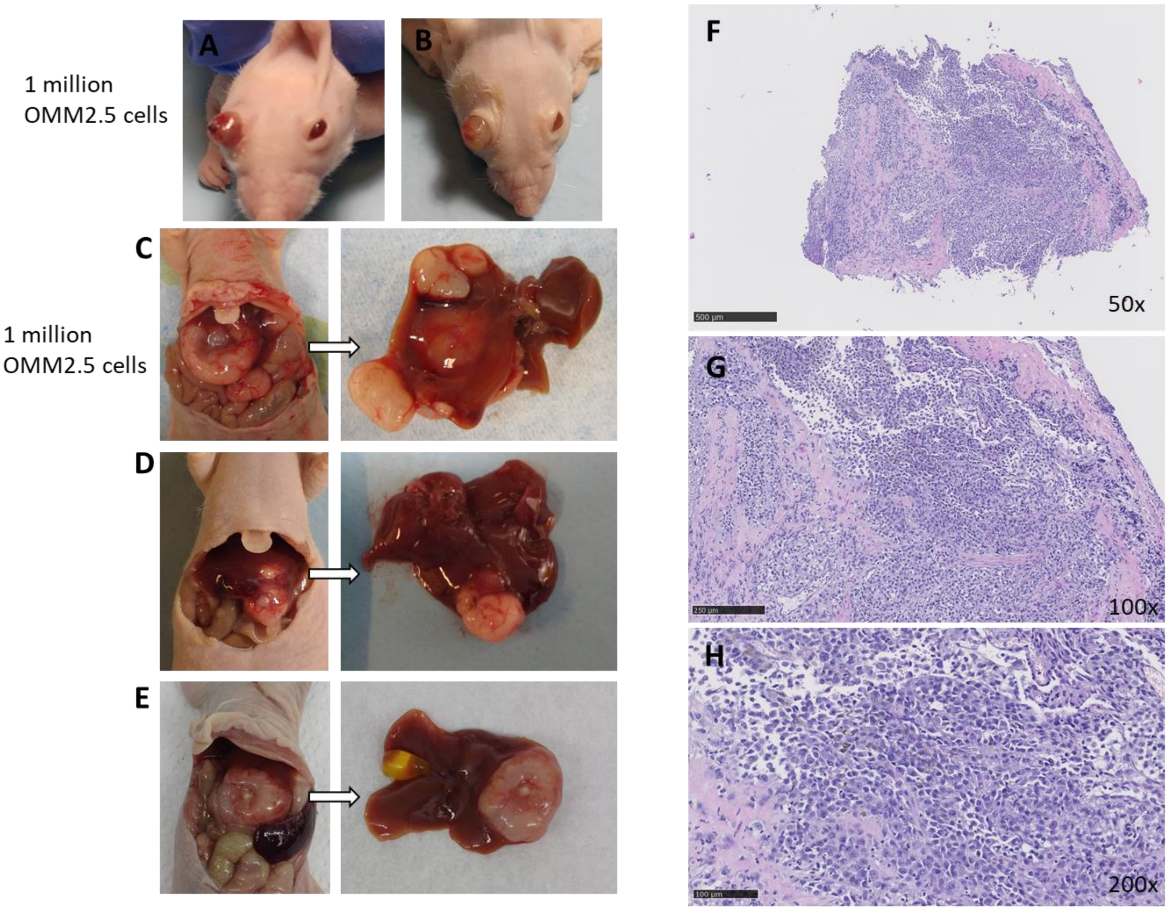


***Supplementary Figure 4* Generation of OMM2.5 cell line-derived orthotopic xenograft models of UM**.

Macroscopic appearance of an ocular tumour (A) 104 days after intraocular injection of 1 million OMM2.5 cells, or (B) 102 days after intraocular injection of disaggregated cells from ocular tumour (A). Macroscopic appearance and liver gross pathology of two representative mice injected with 1 million OMM2.5 cells in the liver, 98 days post-injection (C), 112 days post-injection( D), and of one representative mouse directly implanted with a OMM2.5 cell line-derived tumour fragment in the liver (E), 61 days post-implantation (E). H&E staining in representative sections of an ocular tumour from an OMM2.5 cell line- derived xenograft model. Original magnification x50 (F), x100 (G), x200 (H).
